# Bioenergetic reprogramming of macrophages reduces drug tolerance in *Mycobacterium tuberculosis*

**DOI:** 10.1101/2025.01.24.634842

**Authors:** Vikas Yadav, Sarthak Sahoo, Nitish Malhotra, Richa Mishra, Raju S Rajmani, Siva Shanmugam, Radha K Shandil, Shridhar Narayanan, Vivek V. Thacker, Mohit Jolly, Aswin Sai Narain Seshasayee, Amit Singh

## Abstract

Eradication of *Mycobacterium tuberculosis* (*Mtb*) requires strategies targeting bacteria inside the host. *Mtb* exhibits heterogeneity in redox metabolism inside macrophages to evade killing by anti-TB drugs. If and how macrophage physiology correlates with bacterial redox heterogeneity and drug tolerance remains unclear. Using a fluorescent reporter of mycobacterial redox potential, flow sorting, and RNA sequencing of infected macrophages, we characterized transcriptional and metabolic responses of macrophages harboring redox-diverse *Mtb* populations. We found that macrophages with suppressed glycolysis and elevated oxidative phosphorylation (OXPHOS) correlated with *Mtb* populations exhibiting reductive stress and drug tolerance. Conversely, macrophages with elevated glycolysis and suppressed OXPHOS displayed higher mitochondrial reactive oxygen species through reverse electron transport, resulting in oxidative stress in *Mtb* and enhancing drug efficacy. Computational and genetic approaches identified Nrf2 as a key regulator of macrophage bioenergetics driving redox heterogeneity and drug tolerance in *Mtb*. Redirecting macrophage metabolism from OXPHOS to glycolysis using an FDA-approved antiemetic drug, meclizine, subverted redox heterogeneity and diminished drug tolerance in macrophages and mice. The pharmacological profile of meclizine (C_max_ and AUC_last_) indicated no adverse interactions with first-line anti-TB drugs in mice. Our data demonstrate the feasibility of reprogramming macrophage metabolism to reduce drug tolerance in *Mtb* infection.

## Introduction

Infection with the human pathogen *Mycobacterium tuberculosis* (*Mtb*) represents one of the most intricate inter-organismic interactions in which both pathogen and host cells (e.g., macrophages) exhibit remarkable phenotypic diversity [1]. Examples of phenotypic heterogeneity within infected macrophages include variations in polarization states (M1/M2), carbon metabolism (glycolysis/fatty acid oxidation), and ontogeny (alveolar macrophage [AMs]/interstitial macrophage [IMs] lineages). This heterogeneity is one of the critical determinants that contribute to the need for prolonged treatment of tuberculosis (TB), which results in noncompliance and the emergence of drug-resistant infections [2, 3]. Our mechanistic understanding of how heterogenous host-pathogen interaction dictates the outcome of antimycobacterial therapy is incomplete: filling this knowledge gap is essential for shortening lengthy TB treatment regimens and finding new ways to support future drug discovery programs.

*In vivo* studies show that diverse functional states of macrophages provide different niches for *Mtb* that distinctly affect bacterial physiology and the ability not to be killed by drugs [4–7]. IFNγ-mediated activation of macrophages induces phenotypic drug tolerance by shifting *Mtb* from an actively proliferating to a non-proliferating metabolically active state [4]. A genome-wide screen using *Mtb-*infected macrophages reported a role for magnesium transporter (MMGT1) and lipid droplets in switching *Mtb* from an actively replicating to a stressed, non-replicating state of persistence [8]. The uniform linkage between diminished growth rate and drug tolerance is reinforced by studies showing that nitric oxide (NO) produced by macrophages inhibits bacterial respiration, Fe-S cluster homeostasis, and central carbon metabolism, thereby inducing metabolic quiescence and phenotypic drug tolerance [4, 9, 10]. Notably, the growth rate of *Mtb* varies significantly among individual macrophages and correlates well with the pre-existing cell-cell variation of inducible nitric oxide synthase (iNOS) activity [11]. Similarly, an extreme form of drug tolerance is displayed by non-replicating *Mtb* present in the cavity caseum obtained from infected rabbits [12]. Clinically prevalent mutations in genes involved in carbon metabolism (glycerol (*glpK*), acetate (*icl1*) and propionate (*prpR*)) and toxin-antitoxin modules cause fitness defects and mediate multi-drug tolerance in *Mtb* inside macrophages and animals, possibly undermining antibiotic efficacy in humans [13–18].

While stress-induced metabolic quiescence and increase in drug tolerance are frequently reported in *Mtb* [4, 19], drug tolerance is also exhibited by actively replicating *Mtb* inside macrophages [20, 21]. Animal experiments show that a subpopulation of replicating and non-replicating bacteria resume growth after drug withdrawal [19, 20], indicating that growth-arrested bacteria are not the sole contributor to tolerance. Consistent with this, a recent study using alveolar macrophages derived from lung lesions of pulmonary TB patients confirmed the presence of both metabolically active and quiescent subpopulations showing multi-drug tolerance [22]. These findings reinforce the idea that mechanistic dissection of host and bacterial determinants associated with drug tolerance in metabolically active *Mtb* needs careful attention to understand why protracted TB therapy is required.

We previously found a functional link between redox heterogeneity and drug tolerance in replicating *Mtb* [21, 23]. By using a ratiometric biosensor (Mrx1-roGFP2) that measures the redox potential of the major cytoplasmic antioxidant mycothiol (*E*_MSH_), we showed that replicating *Mtb* displays redox heterogeneity inside macrophages [21, 23]. Importantly, redox diversity in bacterial populations originates from heterogeneity in the invaded macrophage population. Three distinct macrophage fractions were observed: predominantly harboring *Mtb* in *E*_MSH_-reduced (-300 mV), *E*_MSH_-basal (-275 mV), or *E*_MSH_-oxidized (>-240 mV) *Mtb* states. The host and *Mtb* heterogeneity resulted in differential susceptibilities to anti-TB drugs, with replicating *E*_MSH_-reduced bacteria exhibiting drug tolerance. In contrast, *E*_MSH_-basal bacteria showed intermediate and *E*_MSH_-oxidized bacteria demonstrated extreme sensitivity to anti-TB drugs [23]. Despite the importance of heterogeneity, which physiological aspect(s) contributes to macrophage heterogeneity and how it translates into redox diversification and drug tolerance in replicating *Mtb* remains unexplored. Furthermore, the question of whether the observed heterogeneity in the macrophage population is a cause or consequence of redox diversity in intra-phagosomal *Mtb* remains unresolved.

In the present study, we functionally dissected the relationship between phenotypic heterogeneity in macrophages and redox diversity in *the Mtb* population during infection. Using the redox biosensor combined with flow sorting and RNA sequencing, we uncovered the mechanistic basis of macrophage heterogeneity that induces redox diversity in *Mtb* populations. The study suggests that differences in gene expression among infected macrophages create metabolically diverse environments, which generate *Mtb* subpopulations with varied redox physiology and drug susceptibility. We applied several computational approaches to multidimensional data and found that a cell-protective antioxidant transcriptional program, mobilized by the master regulator nuclear factor erythroid 2 -related factor 2 (NRF2), effectuates bioenergetic changes in infected macrophages that lead to redox heterogeneity and drug tolerance in the *Mtb* population. Genetic and pharmacological reprogramming of mitochondrial bioenergetics collapsed redox heterogeneity and diminished drug tolerance in *Mtb* inside the infected macrophages and in mice.

## Results

### Transcriptional profiling of macrophage sub-populations identifies factors that promote redox diversity and drug tolerance in intracellular *Mtb*

We infected primary mouse bone marrow-derived macrophages (BMDMs) with an *Mtb* strain carrying an established redox biosensor (*Mtb*-roGFP2) plasmid that reports *E*_MSH_ of *Mtb* [23]. Biosensor expression leads to ratiometric changes in the fluorescence excitation at 405 and 488 nm with a uniform emission at 510 nm in response to redox changes in *Mtb*. Under oxidative and reductive conditions, the fluorescence ratio of 405/488 increases and decreases, respectively [23]. We used our previously established flow sorting pipeline that averages the median fluorescence ratio (405/488) of the biosensor expressed by intraphagosomal *Mtb*-roGFP2 to the gate and sort BMDMs into subsets enriched with either *E*_MSH_-reduced, *E*_MSH_-basal, or *E*_MSH_-oxidized bacteria (Fig S1). We previously showed that *E*_MSH_-reduced bacteria inside THP-1 macrophages are replicative and tolerant to isoniazid (INH) and rifampicin (RIF), whereas *E*_MSH_-basal and -oxidized *Mtb* showed drug sensitivity [21, 23]. We first determined whether the observations made with the THP-1 cell line and first-line anti-TB drugs are recapitulated in the present experimental setup of primary cells (BMDMs) exposed to multiple antibiotics clinically used to treat drug-sensitive and -resistant TB. Using the biosensor-based gating strategy, we sorted BMDMs infected with *Mtb*-roGFP2 at 24 h post-infection (p.i,), treated them with anti-TB drugs (isoniazid, rifampicin, moxifloxacin, and bedaquiline at 3X the *in vitro* minimum inhibitory concentration (MIC)) for 48 h, and confirmed that the *E*_MSH_-reduced fraction is uniformly more tolerant to multiple anti-TB drugs than the *E*_MSH_-oxidized fraction (Fig. S2). In subsequent experiments, we focussed on BMDMs harboring *E*_MSH_-reduced or *E*_MSH_-oxidized bacteria, as these fractions represent drug-tolerant or drug-sensitive phenotypes of *Mtb*, respectively [21, 23].

To acquire a mechanistic understanding of why macrophage subsets harbor redox diverse *Mtb* populations, we sorted *Mtb*-roGFP2-infected BMDMs 24 h p.i. and subjected them to RNA-sequencing (RNA-seq) (Fig. 1A). Four macrophage fractions were sorted for analysis by RNA-seq: BMDMs-predominantly harboring (i) *E*_MSH_- reduced bacilli and, (ii) *E*_MSH_-oxidized bacilli, (iii) bystander BMDMs, and (iv) uninfected (Fig. 1A). We identified differentially expressed genes (DEG) [base mean > 10, fold change > 1.5, false discovery rate (FDR)< 0.1] between *Mtb*-infected BMDM subsets, bystanders, and uninfected BMDMs (Table S1). Principal components analysis (PCA) showed that samples clustered with their biological replicates with clear segregation between uninfected and *Mtb*-infected BMDMs (Fig. 1B). An additional PCA on all infected BMDMs consistently revealed clear segregation of two populations of cells containing drug-tolerant (*E*_MSH_-reduced) or -sensitive (*E*_MSH_- oxidized) *Mtb*, indicating differences in the transcriptome signatures (Fig. 1C). Compared to the uninfected control, the expression of 6460 and 6225 genes was affected in the BMDMs harboring *E*_MSH_-reduced and *E*_MSH_-oxidized populations, respectively (Fig. 1D and Table S1). Direct comparison of transcriptomes revealed that 825 genes were differentially regulated between *E*_MSH_-oxidized and *E*_MSH_-reduced fractions (374 and 367 genes were upregulated in *E*_MSH_-oxidized and *E*_MSH_-reduced fractions, respectively) (Fig. 1D and Table S2). Expression of 6005 genes was deregulated between bystanders and uninfected BMDMs (Table S1). The bystander transcriptome overlapped with both the infected subpopulations (fold change > 2.0, false discovery rate [FDR]< 0.1); the extent of overlap was ∼ 2-fold more with the *E*_MSH_- oxidized fraction (160 vs. 84 genes, Fig S3A Table S3). The uniquely overlapping genes between bystander BMDMs and *E*_MSH_-oxidized BMDMs were 212 compared to 99 for the bystander vs *E*_MSH_-reduced fractions (Fig S3A). Consistent with this result, flow sorting of bystander BMDMs and subsequent infection with *Mtb*-roGFP2 uniformly shifted the *E*_MSH_ of *Mtb* towards the oxidative state (Fig. S6).

**Fig. 1.**
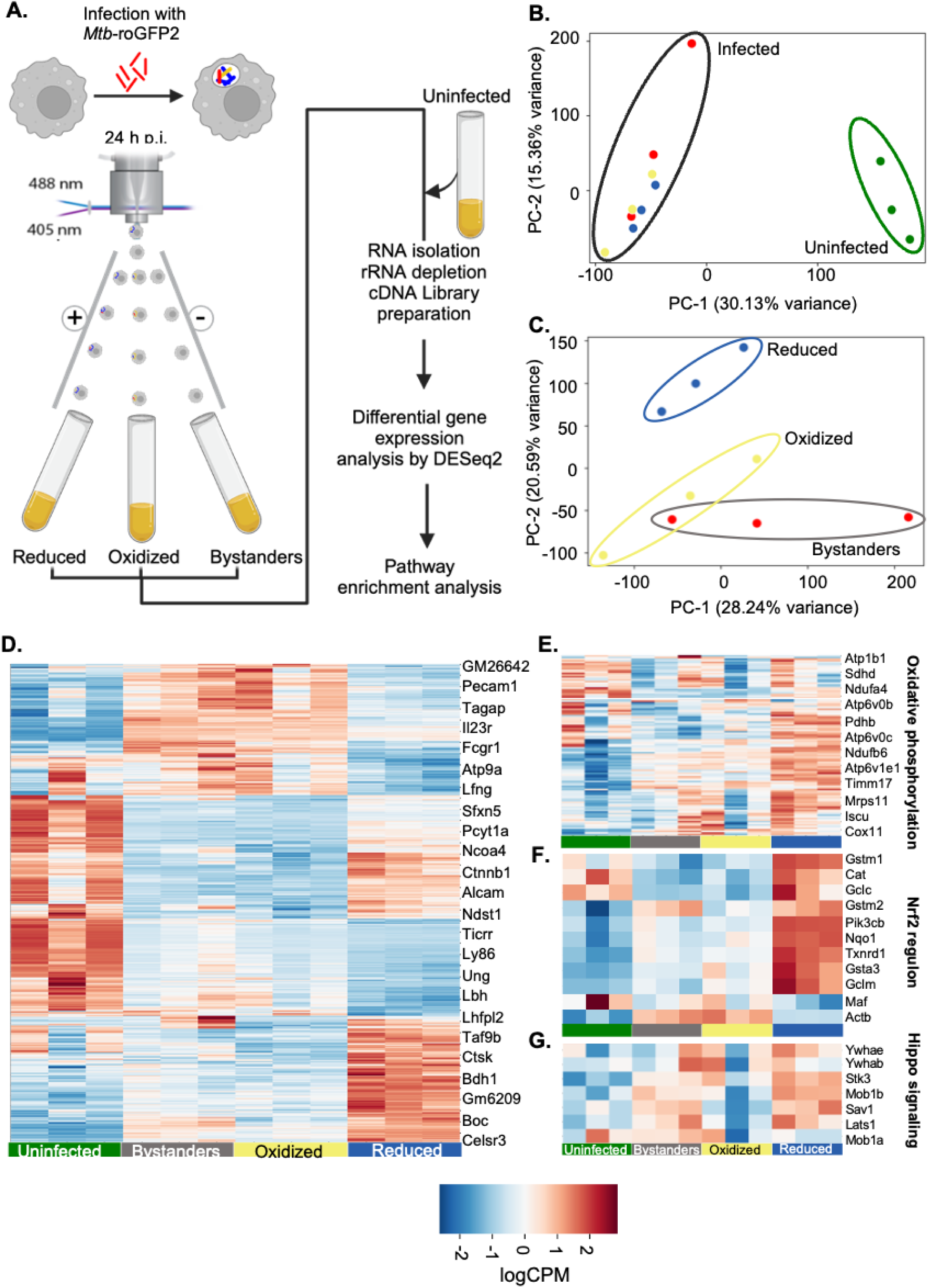
Transcriptomic profiling of macrophages harboring redox diverse *Mtb*. **A.** Schematic showing flow sorting–coupled RNA sequencing of macrophage subpopulations: uninfected (green), bystanders (grey), macrophages harboring *E*_MSH_- oxidized (yellow, here on labeled as ‘oxidized’) or *E*_MSH_-reduced (blue, here on labeled as ‘reduced’) *Mtb*.; **B.** Principal component analysis (PCA) plot comparing infected and uninfected macrophages; **C.** PCA plot showing the subpopulations within the infected macrophages: ‘bystanders’, ‘oxidized’, ‘reduced’; **D.** Heat map of genes differentially expressed between the subpopulations; genes clustered according to their involvement in pathways: **E.** Oxidative phosphorylation. **F.** Nrf2 regulon. **G.** Hippo signaling. Heatmaps are generated from three independent experiments with base mean >10, FDR < 0.1, and log2FC > 0.6.

The transcriptome of *E*_MSH_-oxidised and *E*_MSH_-reduced BMDMs overlapped considerably with that of previously reported transcriptional data of diverse macrophage populations infected with *Mtb ex vivo* and *in vivo* (BMDMs, AM, and IM) (Fig. S3B-D) [5, 24]. Overlap analysis between 825 DEGs of oxidized vs reduced fractions with the DEGs between *Mtb*-infected AM or IM showed a marginally higher overlap of *E*_MSH_-oxidized BMDMs with IM (17%, Fisher exact test: *p*<0.0001) compared to the *E*_MSH_-reduced BMDMs (11%, Fisher exact test: *p*<0.0001) (Fig S3B and Table S4). DEGs between oxidized and reduced fractions significantly overlapped with *Mtb-*infected AMs isolated from the bronchoalveolar lavage (BAL) fluid of mice during the early stages of infection (day 10 p.i.) (Fig S3C and Table S5) [25], confirming the uniformity in the primary response of macrophages to infection *ex vivo* and *in vivo*. However, among the overlapping genes, the early transcriptome of bronchoalveolar lavage (BAL) fluid-isolated murine AMs showed higher similarity with BMDMs containing *E*_MSH_-reduced than *E*_MSH_-oxidized bacteria (47 vs 15 respectively; Fig S3C), suggesting the macrophages harboring *E*_MSH_-reduced *Mtb* are AM-like in their transcriptional response. We did not find any significant overlap with the transcriptomes of M1 and M2 macrophages (Fig S3D and Table S6), which suggests that these macrophage subpopulations do not segregate along the M1-M2 axis.

To understand the basis of drug tolerance exhibited by *E*_MSH_-reduced bacilli in BMDMs (Fig. S2), we performed pathway analysis using the gene ontology tool (ShinyGO 0.80 [26]) and pathway enrichment tool (Enrichr [27]). We found that genes associated with mitochondrial respiration (oxidative phosphorylation [OXPHOS]), central carbon metabolism (TCA cycle), Hippo signaling, and cell protective antioxidant responses (Nrf2- regulon) were induced more in BMDMs containing *E*_MSH_-reduced bacilli than in -oxidized bacilli (Fig. 1D-G). NRF2 is a master regulator of a cell-protective antioxidant transcriptional signature, including antioxidant production (Nqo1, Cat, Txnrd1), iron metabolism (Slc7a11, Slc7a2, Clec4e, Fbxl5), and cytoplasmic thiol production (*Gsta3, Gclc, Gstm1* and *Gstm2*). These genes were induced in macrophages with *E*_MSH_-reduced fraction versus -oxidized fraction (Fig. 1F, Table S2). Consistent with this, using a transcription factor enrichment tool (ChEA3 [28]), we found that the genes upregulated in BMDMs harboring *E*_MSH_-reduced bacteria contained an Nrf2-binding motif (Fig S4A). Enrichment of Nrf2-specific genes was further substantiated by a significant commonality (enrichment FDR: 1.0E-05) with the genes downregulated in the transcriptome of lung tissues of Nrf2 knock-out (Nrf2_-/-_) transgenic mice (Fig. S4B). Another transcription factor, Bach2, negatively regulates the antioxidant response by suppressing Nrf2 transcriptional activity [29, 30]. As expected, genes upregulated in lung macrophages of Bach2 knock-out (Bach2^-/-^) mice overlapped significantly with genes induced in BMDMs harboring *E*_MSH_-reduced bacteria (enrichment FDR: 3.0E-24, Fig S4B). Ontology-based pathway enrichment further established the upregulation of antioxidant pathways in the macrophages harboring *E*_MSH_-reduced *Mtb* (Fig. S4C). The oxidized fraction of infected BMDMs showed enrichment of genes involved in the cell cycle, DNA damage response (base-excision, DNA mismatch repair), and complement activation pathways (Fig S5).

Activation of the Nrf2 and Hippo pathways, which respond to redox and energetic changes [31–36], along with upregulation of the TCA cycle and OXPHOS genes in BMDMs containing *E*_MSH_-reduced bacteria, suggest that host energy homeostasis could be an important physiological parameter required for mediating redox-dependent drug tolerance in *Mtb* populations.

### Nrf2-dependent changes in mitochondrial bioenergetics induce redox-linked drug tolerance in *Mtb*

Our transcriptomic data suggest a role for mechanisms controlling host redox balance and mitochondrial bioenergetics in supporting the emergence of a drug-tolerant *E*_MSH_- reduced population during infection. To clarify this link, we sorted BMDMs enriched with *E*_MSH_-reduced or *E*_MSH_-oxidized bacteria and applied an extracellular flux analyzer to measure oxygen consumption rate (OCR) and extracellular acidification rate (ECAR) (Fig. 2A). Mammalian cells rely on cytosolic glycolysis and mitochondrial oxidative phosphorylation (OXPHOS) to generate energy for cellular functions. The OCR quantifies OXPHOS, whereas ECAR quantifies acidification due to lactate export during glycolytic flux [37]. Consistent with our RNA-seq data, BMDMs harboring *E*_MSH_- reduced *Mtb* exhibited higher basal respiration and ATP-linked OCR than *E*_MSH_- oxidized fraction (Fig. 2B and C). Conversely, glycolytic parameters, such as glucose metabolism and glycolytic capacities, were significantly higher in *E*_MSH_-oxidized BMDMs than in the *E*_MSH_-reduced fraction (Fig. 2D and E).

**Fig 2:**
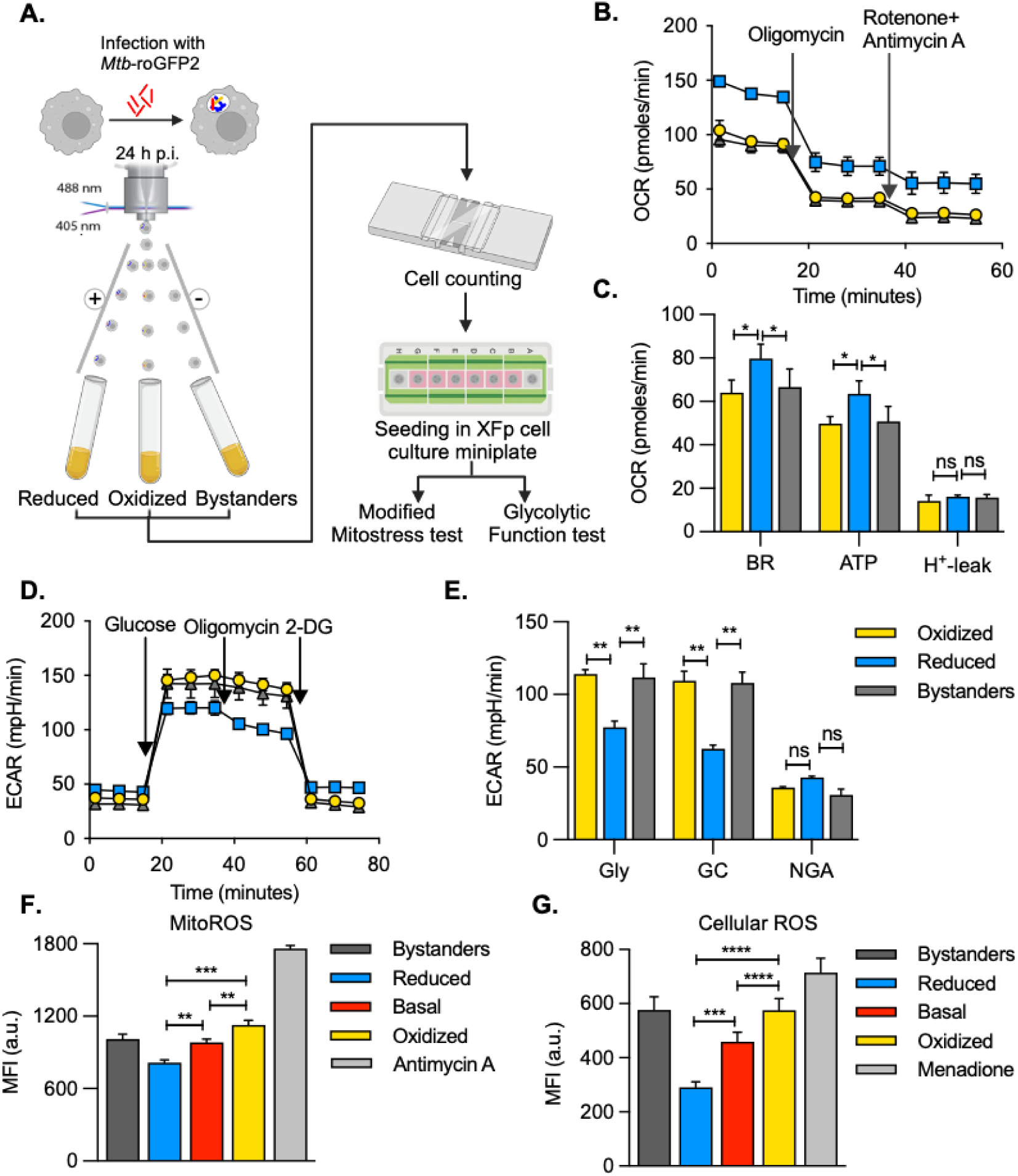
Bioenergetically heterogenous BMDMs harbor redox-diverse bacteria. **A.** Workflow of the flow-sorting coupled seahorse extracellular flux analysis; **B. & C.** A modified mitostress test was performed to calculate mitochondrial parameters. BR- basal respiration, ATP- ATP production, H+-leak- proton leak; **D. & E.** ECAR test was performed to assess the parameters associated with glycolysis in the three sorted BMDM subpopulations. Gly- glycolysis, GC- glycolytic capacity, NGA- non-glycolytic acidification; **F.** Mitochondrial ROS in *Mtb*-roGFP2 infected BMDMs. Antimycin A used as the positive control; **G.** Cellular ROS measured in *Mtb*-roGFP2 infected BMDMs, 100 µM menadione used as the control. MFI- median fluorescence intensity, a.u.- arbitrary units. Data are expressed as mean ± S.D representative of three independent experiments. *p*-value determined using an unpaired t-test with Welch’s correction. ns- non-significant, \**p*<0.05, \*\**p*<0.01.

The drop in mitochondrial OXPHOS in BMDMs containing *E*_MSH_-oxidized bacilli is accompanied by higher mitochondrial reactive oxygen species (mitoROS, a hallmark of mitochondrial stress) compared to BMDMs harboring *E*_MSH_-reduced bacteria (Fig. 2F). MitoROS is a significant contributor to cellular ROS [38]. In agreement with this, the accumulation of cellular ROS was ∼3-fold higher in BMDMs containing *E*_MSH_- oxidized *Mtb* than in the *E*_MSH_-reduced fraction (Fig. 2G). The generation of ROS is dependent on the electron transport chain (ETC) during forward electron transport (FET), as well as when electrons flow backwards (reverse electron transport or RET) through complex I [39]. To establish the source of ROS (FET or RET) within the ETC of BMDMs, we measured mitoROS in BMDMs treated with rotenone. In FET, rotenone facilitates ROS generation at complex I. In contrast, rotenone reduces ROS by blocking complex I during RET [39, 40]. The addition of rotenone uniformly reduces mitoROS in all the subsets of BMDMs infected with *Mtb*, consistent with earlier reports of RET as the primary contributor of mitoROS during *Mtb* infection [40] (Fig. S7). Consistent with the RNA-seq data, the bystander BMDMs exhibit respiratory and redox changes similar to BMDMs containing *E*_MSH_-oxidized *Mtb* (Fig. 2A-G). These data demonstrate that BMDMs containing drug-tolerant *E*_MSH_-reduced bacilli maintain mitochondrial function but restrained glycolytic flux, whereas the *E*_MSH_-oxidized fraction of BMDMs experience glycolytic shift and mitochondrial stress.

Nrf2 is a prominent factor that sustains the structural and functional integrity of mitochondria (*e.g.,* ATP synthesis, membrane potential, mitoROS, and substrate availability) [41, 42] and that is overexpressed in alveolar macrophages during the early stages of *Mtb* replication in mice [25]. Since an Nrf2-driven transcriptional response featured in our RNA-Seq analysis, we hypothesized that Nrf2 could link macrophage bioenergetics to redox diversity in *Mtb*. Supporting this, we found that several genes upregulated in macrophages containing *E*_MSH_-reduced *Mtb* had an NRF2 ChIP-Seq peak [25, 43]. In contrast, none of the genes upregulated in the macrophages harboring *E*_MSH_-oxidized *Mtb* showed these peaks (Fig. 3A). On this basis, we reduced the expression of Nrf2 in BMDMs using siRNA (siNRF2-BMDMs), resulting in a significant down-regulation of Nrf2 and its dependent genes (Fig. S8A). As expected, the knockdown of Nrf2 increased mitoROS more than the scrambled siRNA (siSCR-BMDMs) control upon *Mtb* infection (Fig. 3B). We next investigated the effect of Nrf2 knockdown on mitochondrial respiration, *E*_MSH_ of *Mtb*, and INH tolerance. Basal OCR and ATP-linked OCR were dramatically reduced in siNRF2-BMDMs compared to siSCR-BMDMs at 24 h p.i., indicating decreased mitochondria respiration (Fig. 3C and D). Nrf2 knockdown did not increase glycolysis to compensate for the decreased mitochondrial respiration, which could be due to partial silencing of Nrf2 (Fig. S8B). Diminished mitochondrial respiration and enhanced mitoROS correlated with an increased fraction of siNRF2-BMDMs harboring *E*_MSH_-oxidized *Mtb* relative to siSCR-BMDMs 24 h p.i.(Fig. 3E). We also determined whether the more significant fraction of siNrf2-BMDMs harboring *E*_MSH_-oxidized *Mtb* reduces INH tolerance: survival of *Mtb* was comparable in siNRF2 and siSCR BMDMs at 72 h p.i. We observed that *Mtb* growing in siNRF2 BMDMs were significantly more sensitive to INH than those in siSCR BMDMs (percentage survival in siSCR: 39 % ± 3 %, siNRF2: 19 % ± 1 %) (Fig. 3F).

**Fig. 3.**
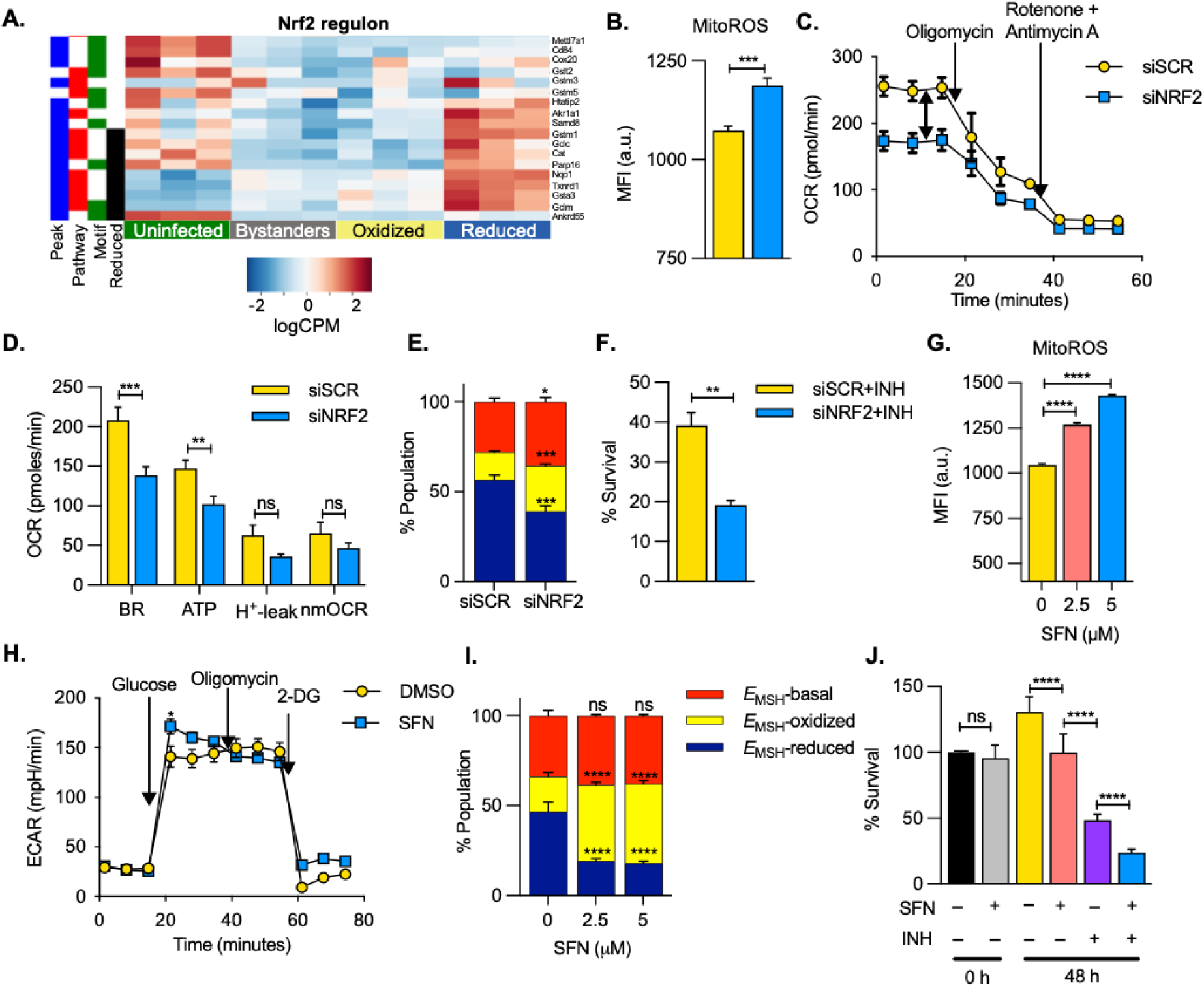
Nrf2 knockdown diminishes *E*_MSH_-reduced fraction and reverses antibiotic tolerance in macrophages. **A.** z-normalized RNA-Seq data of uninfected, bystanders, oxidized, and reduced populations where genes are clustered by expression levels. Only those genes are represented that satisfy two of the following criteria: belong to the Nrf2 pathway (red) and bear the Nrf2 binding motif in the promoter region (green) or have an Nrf2 ChIP-seq (Chromatin immunoprecipitation followed by sequencing) peak detected near the promoter region (blue). Genes in black represent those that are significantly upregulated in the reduced subpopulation in comparison to the oxidized subpopulation (no gene upregulated in the oxidized subpopulation was detected amongst the above genes); **B.** Mitochondrial ROS in BMDMs transfected either with scrambled siRNA (siSCR, yellow) or siRNA targeted against NRF2 (siNRF2, blue) at 24 h p.i. MFI- median fluorescence intensity, a.u.- arbitrary units; **C and D.** Modified mitostress test of siSCR or siNRF2 BMDMs at 24 h p.i. infected with *Mtb* H37Rv at a moi of 2. BR- basal respiration, ATP- ATP production, H+-leak- proton leak, nmOCR- non-mitochondrial respiration; **E.** Redox profile of *Mtb*- roGFP2 infected siSCR or siNRF2 BMDMs at 24 h p.i; **F.** Bar plot showing the percentage survival of *Mtb* in siSCR and siNRF2 BMDMs, treated with 3X MIC of INH (0.375 µg/ml) for 48 h. INH treatment was initiated at 24 h p.i. Percentage survival is calculated compared to the untreated BMDMs; **G.** Mitochondrial ROS assessed in infected BMDMs upon treatment SFN at the indicated concentrations at 24 h p.i.; **H.** Glycolytic function test showing the extracellular acidification rate (ECAR) upon treatment with 5 µM SFN at 24 h p.i.; **I.** Percentage distribution of redox-diverse fractions of *Mtb*-roGFP2 in BMDMs treated with the indicated concentrations of SFN at 24 h p.i.; **J.** Antibiotic tolerance of intracellular *Mtb* assessed by CFUs. Percentage survival is compared to 0 h untreated group. Data are expressed as mean ± S.D. of three independent experiments. *p*-value determined using an unpaired t-test with Welch’s correction. ns- non-significant, \**p*<0.05, \*\**p*<0.01, \*\*\**p*<0.001, \*\*\*\**p*<0.0001.

As an additional verification of a functional linkage between Nrf2 and mycobacterial redox, we used sorafenib, a pharmacological inhibitor of the Nrf2-Keap1 axis [44]. Sorafenib attenuates the nuclear translocation of Nrf2 and suppresses the transcriptional expression of Nrf2-dependent antioxidant and respiratory genes, resulting in elevated mitochondrial stress [44, 45]. Consistent with this observation, BMDMs treated with non-toxic concentrations of sorafenib (SFN, 2.5-5 μM) led to suppressed OCR, a compensatory increase in glycolytic rate, and enhanced mitoROS (Fig. 3G-H, S9A). Importantly, SFN treatment of BMDMs diminished redox heterogeneity in intra-phagosomal *Mtb,* with most of the BMDMs harboring bacteria in the *E*_MSH_-basal or -oxidized states (Fig. 3I). Lastly, 50% of *Mtb* survived INH treatment in BMDMs, whereas survival of *Mtb* reduced to 24% upon exposure to SFN (5 μM) with INH (Fig. 3J). We confirmed that 5 μM of SFN does not affect *Mtb* survival in the 7H9 growth medium, macrophage viability, and the MIC of SFN against *Mtb* is 20 μM (Fig. S9B-D), suggesting that SFN lowers INH tolerance of intra-phagosomal *Mtb* by modulating host pathways. These data indicate that the differential induction of Nrf2 is one of the mechanisms that account for the observed bioenergetic differences in macrophages, resulting in redox heterogeneity and drug tolerance of *Mtb*.

### Metabolic state of macrophages drives redox-dependent drug tolerance in *Mtb*

Data indicate a preferential engagement of mitochondrial OXPHOS by BMDMs harboring drug-tolerant *E*_MSH_-reduced bacteria, whereas drug-sensitive *E*_MSH_-oxidized *Mtb* resides in BMDMs committed to glycolysis. These observations motivated us to determine whether shifting metabolic reliance interchangeably between mitochondrial respiration and glycolysis modulates redox heterogeneity and drug tolerance in *Mtb*. To do this, we first examined the energy source driving mitochondrial respiration and the emergence of drug-tolerant, *E*_MSH_-reduced *Mtb* in BMDMs.

Mitochondria utilize glucose, glutamine (Gln), and fatty acids to generate ATP by OXPHOS. We assessed the preference of BMDMs harboring redox-diverse *Mtb* to use glucose, Gln, or fatty acids as energy sources in the mitochondrial stress test. To test the oxidation of endogenous fatty acids, we flow-sorted BMDMs containing *E*_MSH_- reduced and *E*_MSH_-oxidized *Mtb*. We measured OCR after treatment with the fatty acid oxidation inhibitor etomoxir (Eto). Both basal OCR and ATP-linked OCR were significantly reduced upon Eto treatment of BMDMs containing *E*_MSH_-reduced bacteria (Fig. S10A). However, Eto treatment also suppresses the OCR of BMDMs harboring *E*_MSH_-oxidized and bystander BMDMs (Fig. S10B). Moreover, Eto treatment did not affect the redox heterogeneity displayed by intra-phagosomal *Mtb*. Like Eto, trimetazidine, which inhibits the 3-keto acyl CoA thiolase step in the β-oxidation of fatty acids [38, 46], had no influence on redox heterogeneity exhibited by intra-phagosomal *Mtb* (Fig. S10C). These findings agree with fatty acids being a principal energy source for *Mtb*-infected macrophages [47–49]. However, the fatty acid-dependent changes in the metabolic state of infected macrophages are unlikely to cause redox variations in *the Mtb* population.

Inhibition of Gln oxidation by bis-2-(5-phenylacetamido-1,3,4-thiadia-zol-2-yl)ethyl sulphide (BPTES) did not affect the OCR of bystander or BMDMs containing redox diverse *Mtb* (Fig. S10D). As expected, BPTES treatment did not influence the redox heterogeneity of *Mtb* (Fig. S10E). Inspection of RNA-seq data revealed enhanced expression of mitochondrially located pyruvate dehydrogenase complex (PDC) in BMDMs harboring *E*_MSH_-reduced compared to *E*_MSH_-oxidized fraction (Fig. 4A). We hypothesize that the *E*_MSH_-reduced fraction utilizes pyruvate for mitochondrial OXPHOS rather than lactic acid production. Inhibition of mitochondrial oxidation of glucose upon treatment with UK5099, which blocks pyruvate transport from the cytoplasm into mitochondria, resulted in a significant reduction in basal and ATP-linked OCR of BMDMs (Fig. 4B-D). Importantly, and in contrast to Eto and BPTES, UK5099- mediated inhibition of OCR correlated with a decrease in *E*_MSH_-reduced bacteria with a concomitant increase in *E*_MSH_-oxidized and basal fractions in a concentration-dependent manner (Fig. 4E). Consistent with this, drug tolerance assays suggest that UK5099 uniformly weakens the ability of *Mtb* to tolerate INH and MOXI (Fig. 4F; survival in INH: 32% ± 3%, INH+UK5099: 18% ± 1%; MOXI: 52% ± 3%, MOXI+UK5099: 34% ± 4%).

**Fig. 4:**
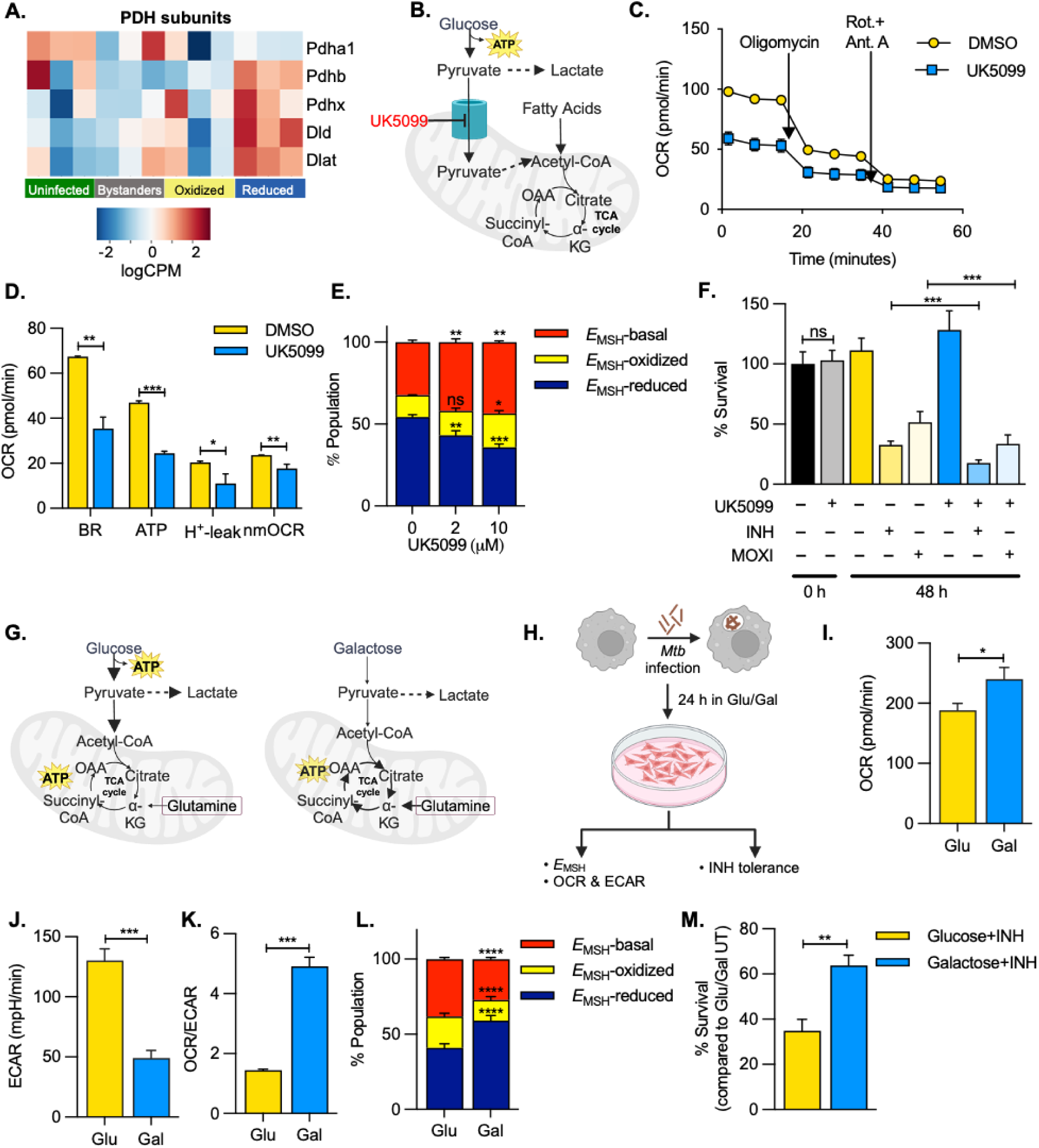
**Macrophage metabolism affects redox physiology and antibiotic tolerance of intracellular *Mtb***. **A.** Heat map showing expression of different subunits of the pyruvate dehydrogenase complex in the macrophage subpopulations: uninfected (green), bystanders (grey), macrophages harboring *E*_MSH_-oxidized (yellow) or *E*_MSH_-reduced (blue) *Mtb;* **B.** Mechanism of action of UK5099, a mitochondrial pyruvate carrier (MPC) inhibitor; **C and D.** Modified mitostress test of *Mtb*-infected BMDMs upon treatment with 10 µM UK5099 at 24 h p.i. BR- basal respiration, ATP- ATP production, H+-leak- proton leak, nmOCR- non-mitochondrial respiration; **E.** Redox profile of intracellular *Mtb* in BMDMs 24h p.i. treated with indicated concentrations of UK5099; **F.** Antibiotic tolerance to 3X MIC of INH (MIC: 0.125 µg/ml) or moxifloxacin (MOXI; MIC: 0.25 µg/ml) with or without 10 µM UK5099. Percentage survival is compared to untreated cells at 0 h p.i.; **G.** Schematic representation of cellular metabolic pathways in the presence of 10 mM glucose or 10 mM galactose as sole sugar sources; **H.** Experimental design to determine the effect of glucose and galactose on the redox poise and antibiotic tolerance of intracellular *Mtb;* **I-K.** Extracellular flux analysis of *Mtb*-infected BMDMs at 24h p.i. to measure mitochondrial respiration (I), glycolysis (J), and a ratio of OCR to ECAR (K); **L.** Redox profile of the intracellular *Mtb* in the presence of 10 mM glucose or 10 mM galactose as sole sugars at 24 h p.i.; **M.** Antibiotic tolerance to 3X MIC of INH in glucose- or galactose-containing medium. Percentage survival is compared to untreated cells after 48 h INH treatment. Data are expressed as mean ± S.D. of three independent experiments. p-value determined using an unpaired t-test with Welch’s correction. ns- non-significant, \**p*<0.05, \*\**p*<0.01, \*\*\**p*<0.001.

Our efforts to confirm the contribution of pyruvate to redox-dependent drug tolerance of *Mtb* is complicated by the fact that multiple mitochondrial enzymes (*e.g.,* glutamate-pyruvate transaminase 2 or alanine aminotransferase 2, serine: pyruvate aminotransferase, L-cysteine to pyruvate conversion pathway) directly contribute to pyruvate flux for OXPHOS without the need for transporting cytosolic pyruvate [50–53]. To circumvent this issue, we shifted cellular energy metabolism by selectively culturing BMDMs on glucose or galactose as the sole sugar source. Production of pyruvate due to glycolysis yields two net ATP, while pyruvate yield, via galactose metabolism, generates no net ATP [54]. Therefore, using galactose as the sole sugar source coerces mammalian cells to generate ATP by utilizing pyruvate to fuel mitochondrial OXPHOS (Fig. 4G and H). In agreement with this idea, when *Mtb*- infected BMDMs were grown in galactose, they exhibited a reduction in ECAR, reflecting decreased glycolysis and increased OCR, resulting in a ∼5-fold increase in the OCR/ECAR ratio compared to glucose as the carbon source (Fig. 4I-K). These data are consistent with a switch to pyruvate oxidation by mitochondria when BMDMs use galactose as an energy source.

We next asked whether a shift in metabolic reliance to galactose-linked mitochondrial OXPHOS induces a reductive shift in the *E*_MSH_ of *Mtb*, leading to increased drug tolerance. Compared to glucose, culturing in galactose resulted in a higher fraction of BMDMs harboring *E*_MSH_-reduced *Mtb* (Fig. 4L). The increased fraction of *E*_MSH_- reduced *Mtb* resulted in 63% of bacteria surviving INH in galactose-grown conditions compared to 30% in glucose-grown conditions (Fig. 4M). Taken together, the data show that inherent metabolic plasticity displayed by macrophages in sugar utilization promotes redox diversity in *Mtb* to tolerate antibiotic pressure. Overall, these results indicate that while both fatty acids and glucose support bioenergetics in infected BMDMs, the flux of glucose via pyruvate for OXPHOS is necessary to induce a reductive shift in the *E*_MSH_ of *Mtb*. In contrast, glycolysis likely plays a prominent role in maintaining bioenergetics in BMDMs containing *E*_MSH_-oxidized *Mtb*.

### Anti-emetic drug meclizine reprograms macrophage bioenergetics during infection

Given that macrophages’ intrinsic ability to shift their reliance on mitochondrial OXPHOS relative to glycolysis, thereby promoting redox-dependent drug tolerance, we hypothesized that targeting such a shift could potentially treat TB. However, pharmacological agents that can safely redirect energy metabolism from OXPHOS to glycolysis without compromising therapeutic value are generally limited and completely absent from the TB field. The deployment of a nutrient-sensitized screening strategy has identified an FDA-approved drug, meclizine, that shifts cellular energy metabolism from mitochondrial respiration to glycolysis in a variety of mammalian cells [54, 55]. Meclizine is available without prescription for the treatment of nausea and vomiting, crosses the blood-brain barrier, and, to our knowledge, has not been explored as a treatment for TB [54]. Based on these findings, we investigated the mechanism and potential therapeutic utility of meclizine in targeting redox-driven drug tolerance in *Mtb*.

We first determined whether meclizine induces a reduction in OCR concomitant with an increase in ECAR in *Mtb-*infected BMDMs. Treatment with 20 μM of meclizine reduced mitochondrial OCR and increased ECAR in *Mtb-*infected BMDMs (Fig. 5A and B). Upon meclizine treatment, we also observed increased mitoROS, and mitochondrial depolarization (Fig. 5C, D). A change in cellular metabolism is often associated with mitochondrial remodeling [56, 57]. We assessed whether meclizine treatment affects the mitochondrial architecture by imaging mitochondria in *Mtb*- infected macrophages treated with meclizine and observed that the surface area and the volume of mitochondria reduce significantly upon treatment (Fig. 5E). These findings suggest increased mitochondrial fragmentation upon meclizine treatment, which could lead to a metabolic shift from OXPHOS towards glycolysis [58, 59].

**Fig. 5.**
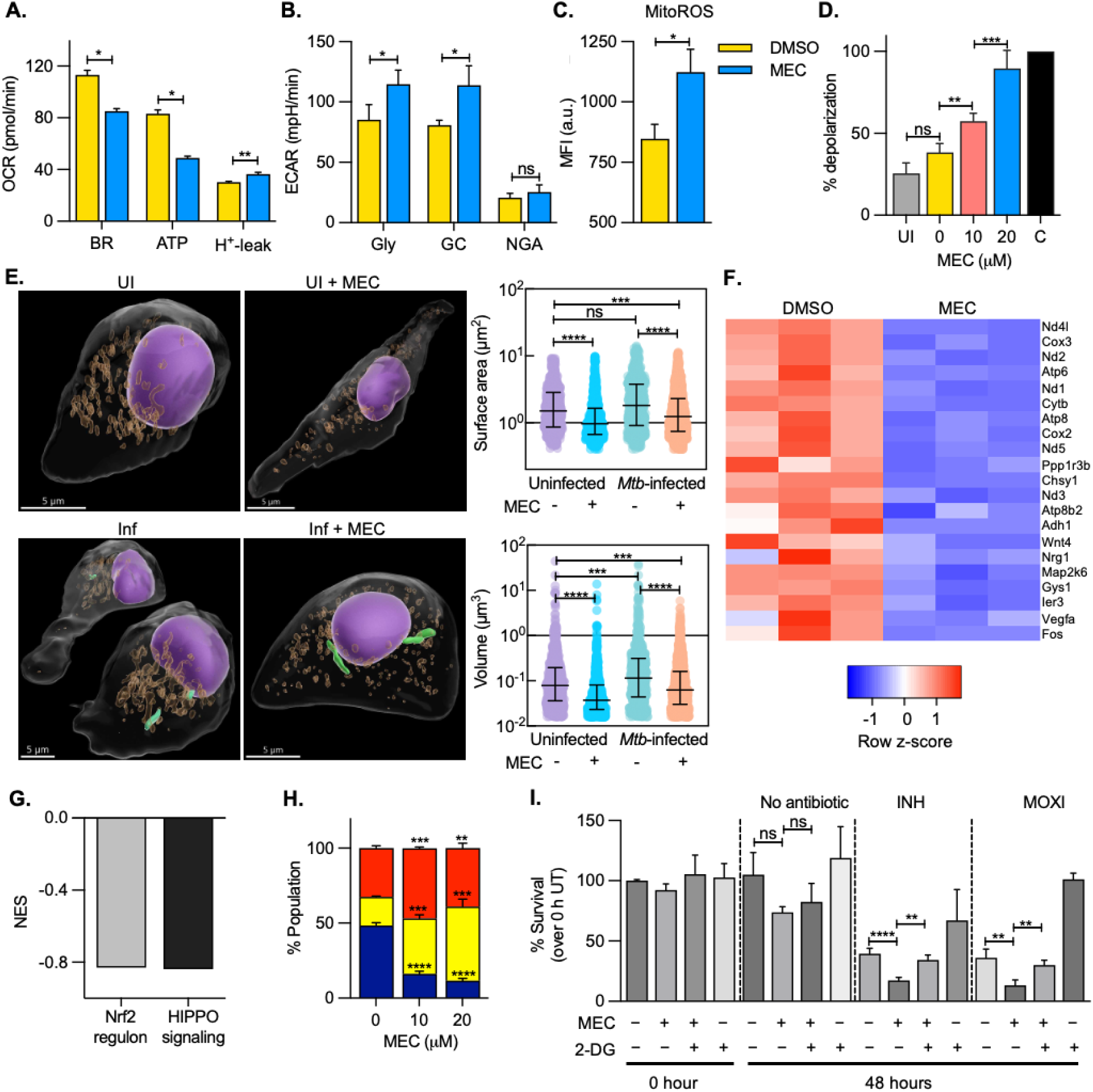
MEC-mediated glycolytic activation diminishes antibiotic tolerance in infected macrophages. **A.** Bar plot showing the mitochondrial respiratory parameters determined by the modified mitostress test at 24 h p.i. upon treatment with 20 µM MEC or 0.2% DMSO (solvent control). BR- basal respiration, ATP- ATP production, H+- leak- proton leak; **B.** Glycolytic stress test to assess the glycolytic levels in infected macrophages 24 h p.i. upon treatment with 20 µM MEC or 0.2% DMSO. Gly- glycolysis, GC- glycolytic capacity, NGA- non-glycolytic acidification; **C.** Mitochondrial ROS measured upon treatment with 20 µM MEC at 24 h p.i.; **D.** Mitochondrial membrane polarization measured by JC-1 dye at 24 h p.i.. 50 µM carbonyl cyanide-p-trifluoromethoxyphenylhydrazone (FCCP) treatment used as positive control; UI: uninfected macrophages; **E.** Representative pseudocoloured 3D views of uninfected and mCherry expressing *Mtb*-infected BMDMs at 24 h p.i., with and without exposure to 20 μM meclizine. Mitochondria (amber), *Mtb* H37Rv (green), and nucleus (purple) are shown as surfaces. Plots show the surface area (µm_2_) and volume (µm_3_) of mitochondria in uninfected and infected BMDMs (n>500 from 6-8 cells per condition). Data are median with interquartile range. p-values calculated using a one-way ANOVA with Dunn’s multiple comparisons test. UI: uninfected BMDMs, Inf: infected BMDMs; **(F. and G.).** RNA-sequencing of BMDMs infected with *Mtb* and treated with 20 µM MEC at 24 h p.i. F. Oxidative phosphorylation gene expression between 0.2% DMSO- treated and MEC-treated BMDMs; and G. net enrichment score (NES) calculated by gene set enrichment analysis (GSEA) for the differentially expressed genes between MEC- and DMSO-treated infected macrophages; **H.** Redox profile of the intracellular *Mtb* in the presence of indicated concentrations of MEC at 24 h p.i.; **I.** Antibiotic tolerance in *Mtb-*BMDMs treated with 20 µM MEC and 0.5 mM 2-DG in the presence of 3X MIC of INH (0.375 µg/ml) or MOXI (0.75 µg/ml). Data are expressed as mean ± S.D. for three biological replicates done in triplicate. p-value determined using an unpaired t-test with Welch’s correction. Ns: non-significant, *p<0.05, **p<0.01, ***p<0.001, ****p<0.0001

However, these changes in mitochondrial function by 20 μM meclizine did not result in the killing of BMDMs infected with *Mtb* (Fig. S11A), likely due to the redirection of metabolism to glycolysis. To interrogate signaling pathways that could explain the metabolic switchover induced by meclizine during infection, we performed global RNA-seq of *Mtb-*infected BMDMs treated with meclizine and untreated control (treated with 0.2% dimethyl sulfoxide [DMSO]) for 24 h. *Mtb*-infected BMDMs treated with meclizine showed differential regulation of 1088 genes compared to the DMSO control (log2-fold change [FC] >0.6, false discovery rate [FDR] < 0.1) (Fig S12A and Table S7). Remarkably, the transcriptional response of meclizine-treated BMDMs reversed the expression of pathways elevated in BMDMs harboring *E*_MSH_-reduced *Mtb* (Fig. 5F and G). For example, meclizine treatment suppressed the Nrf2 regulon, Hippo signaling, and OXPHOS genes (Fig. 5F, G, and S12B). Down-regulation of a significant repertoire of genes associated with OXPHOS explains the suppression of mitochondrial respiration in *Mtb-*infected BMDMs upon meclizine treatment (Fig 5A and F). The lack of deregulation of glycolytic genes suggests that increased glycolysis compensates for diminished mitochondrial respiration. Additionally, meclizine treatment downregulates the expression of the aminoacyl-tRNA biosynthesis pathway (Fig. S12B). One likely possibility is that inhibition of OXPHOS results in ATP limitation, which is critical for tRNA charging. Aminoacyl-tRNA synthetases (ARS) are involved in various cellular processes, such as immune and inflammatory response, angiogenesis, and apoptosis, that can affect *Mtb’s* pathophysiology during infection [60]. Our data indicate that a meclizine-induced metabolic shift is associated with significant changes in the expression of genes controlled by regulators of energy metabolism and redox stress, Nrf2 and Hippo signaling.

### Meclizine collapses redox heterogeneity to diminish drug tolerance in *Mtb*

We then determined whether meclizine-mediated reshaping of macrophage metabolism reverses the drug tolerance and redox heterogeneity displayed by intraphagosomal *Mtb*. Pre-treatment with meclizine abolished the fraction of intra- phagosomal *Mtb* displaying reductive-*E*_MSH_ in a concentration-dependent manner (Fig. 5H). Next, we examined whether meclizine reduced drug tolerance during infection. BMDMs with and without meclizine were infected with *Mtb* for 24 h and exposed to 3X MIC of INH and Mox for an additional 48 h before lysis and enumeration of viable counts. We found that the addition of meclizine reduced INH tolerance significantly (Fig. 5I; INH: 39 % ± 4 %, INH+MEC: 17 % ± 2 %). A similar decrease in tolerance was observed upon substitution of INH with MOXI (MOXI: 36 % ± 7 %, MOXI+MEC: 13 % ± 4 %) (Fig 5I). To determine whether the increased ability of INH and MOXI to kill *Mtb* in macrophages is linked to elevated glycolysis by meclizine, we poisoned glycolysis using an inhibitor of the first hexokinase-mediated step in glycolysis, 2-deoxyglucose (2DG) [5]. Inhibition of glycolysis enhanced bacterial survival in BMDMs and abrogated the potentiating effect of meclizine on lethality induced by INH and MOXI (Fig. 5I). The impact of meclizine (20 μM) is mediated through changes in macrophage metabolism, as extracellular *Mtb* remains viable even at very high meclizine concentrations (320 μM) (Fig S11B and C). Our findings imply that increased OXPHOS and suppressed glycolytic flux in *Mtb*-infected BMDMs induce drug tolerance, which meclizine reverses.

Next, we tested *Mtb’s* response to INH in the presence of meclizine in a murine model of chronic infection. Because the pathophysiology of human TB and tolerance to anti-TB drugs are closely recapitulated in C3HeB/FeJ mice [61], we aerosol-infected this mouse line with *Mtb*. At two weeks p.i, animals were treated with meclizine for two weeks, followed by treatment with INH, meclizine, INH plus meclizine for an additional six weeks, and then we measured the lung bacillary load (Fig 6A). Meclizine dose (25 mg/kg/body weight) was based on previous mouse experiments [62]. As reported earlier, INH monotherapy reduced the bacterial burden from ∼10^6^ to ∼10^4^ per lung at six weeks (*p*=0.0002) of treatment (Fig. 6B). Meclizine alone showed a marginal (∼2.5 fold decrease) effect on bacterial viability over time (Fig. 6B). Relative to the control regimen (INH alone), the addition of meclizine decreased lung CFU by ∼20-fold after six weeks (*p*= 0.0001) of treatment (Fig. 6B). The gross and histopathological changes observed in the lungs after six weeks of therapy were correlated to the observed bacillary load (Fig. 6C and D). The extent of lung damage was greater in the untreated (score = 37.5) and meclizine-treated animals (score = 35), intermediate in INH-treated animals (score = 17.5), and lowest in the case of the meclizine plus INH- treated animals (score = 2.5) (Fig 6E).

**Fig. 6.**
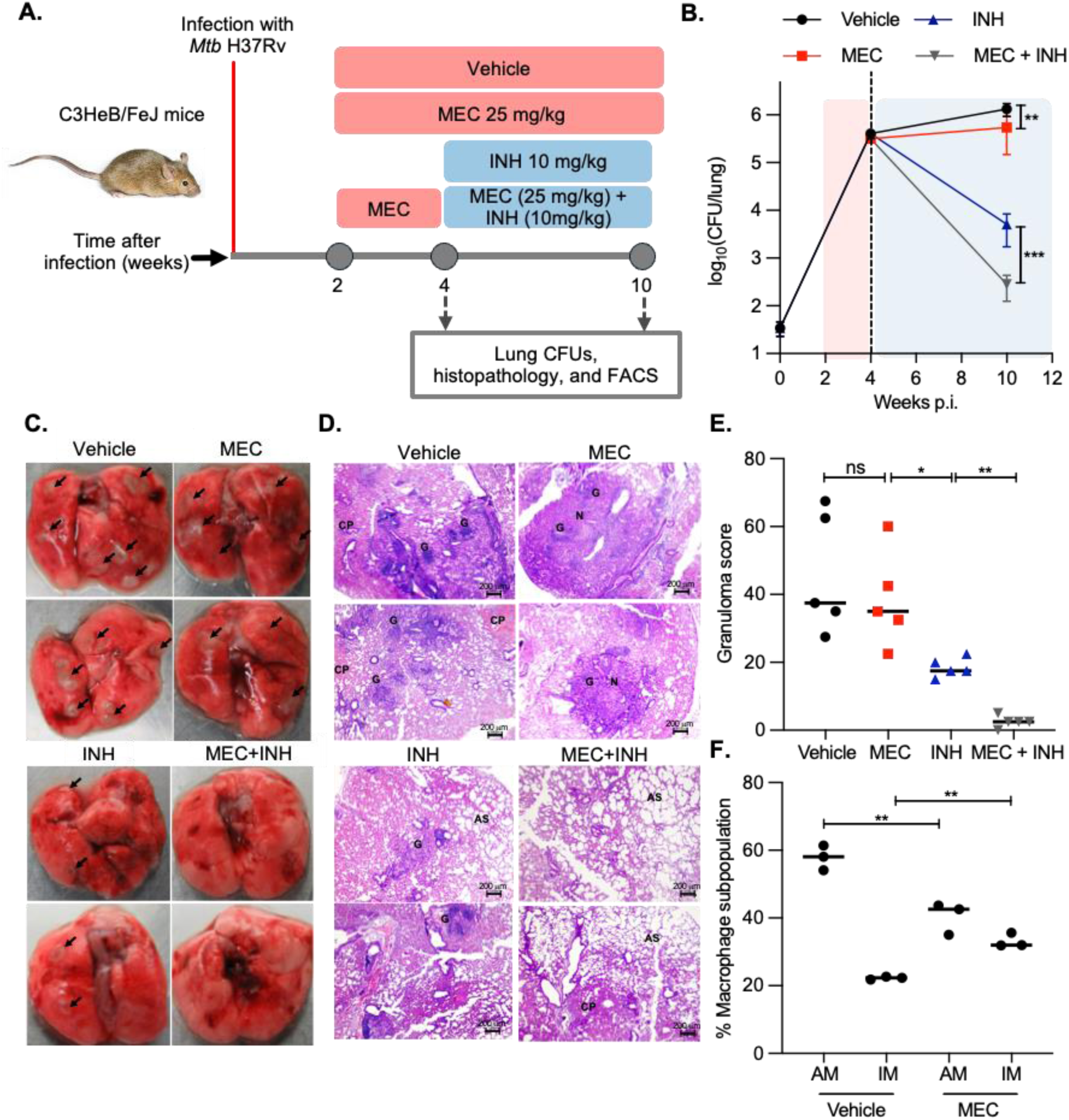
MEC reduces antibiotic tolerance in vivo. **A.** Strategy for investigating the efficacy of MEC at reducing tolerance against INH in C3HeB/FeJ mice; **B.** Bacterial CFUs counted from lungs at the indicated time-points. N≥7 for the 10 weeks p.i. timepoint. Data are expressed as mean ± S.D, and the p-value was determined by the Mann-Whitney test; **C.** Gross pathology of lungs of *Mtb*-infected mice at 10 weeks p.i across experimental groups; **D.** Hematoxylin and eosin–stained lung sections (after 6 weeks of treatment) from mice infected with *Mtb* for all experimental groups. Pathology sections show granuloma (G), alveolar space (AS), collapsed parenchyma (CP), and necrotic area (N). All images were taken at 10X magnification; **E.** Granuloma score was calculated from the histopathological lung sections, **F.** AM and IM populations in the lungs of animals treated with meclizine. For E. and F., data are expressed as mean ± S.D. and the p-value was determined by an unpaired t-test with Welch’s correction. ns-non-significant, \**p*<0.05, \*\**p*<0.01, \*\*\**p*<0.001, \*\*\*\**p*<0.0001.

Since *in vitro* studies indicate that meclizine reduces drug tolerance of intracellular *Mtb* by enhancing glycolysis and suppressing OXPHOS, we also asked whether meclizine stimulates a similar effect *in vivo*. We examined the proportion of alveolar (AM) and interstitial macrophages (IM) upon meclizine treatment. During lung infection in mice, *Mtb* resides in glycolytically active IMs and mitochondrially respiring AMs [5]. Expectedly, treatment of infected mice with the glycolytic inhibitor 2-DG decreased the number of glycolytically active IMs [5]. We then asked whether meclizine alters the proportion of IMs and AMs. In line with *vitro* findings, the proportion of glycolytically active IMs increased, and AMs decreased at six weeks post-meclizine treatment (Fig. 6F). Together, these results confirm that adjunct therapy with meclizine counteracts drug tolerance by reshaping the metabolism of infected macrophages.

### Meclizine exhibits no adverse interaction with anti-TB drugs

A favorable safety profile, oral pharmacokinetics with the ability to cross the blood-brain barrier, and years of clinical use in humans for treating nausea and vertigo [63, 64] make meclizine a good candidate for developing new therapeutic combinations for treating TB. We assessed the pharmacological compatibility of meclizine by measuring its interaction with clinically relevant, first-line anti-TB drugs (isoniazid or H, rifampicin or R, ethambutol or E, and pyrazinamide or Z) given as a combination. We also performed a single-dose pharmacokinetic interaction by administering HREZ orally at the human equivalent doses with and without meclizine (25 mg/kg/body weight, intraperitoneally [i.p]) in mice. We have included an additional group of mice that was dosed with meclizine (25 mg/kg/body weight, intraperitoneally [i.p]) to compare the PK profile of meclizine in the presence of an HREZ combination (Fig. 7A). Plasma and lung homogenates were analyzed for individual drugs using liquid chromatography-mass spectrometry, and parameters such as maximum plasma concentration (C_max_) and area under the plasma concentration-time curve (AUC_last_) were quantified as a ratio for single-treatment groups vs. the combination (Fig. 7B-F). PK profiles revealed no to moderate drug-drug interaction when meclizine was administered with HREZ. C_max_ and AUC_last_ for meclizine, when administered alone, were 0.31 ug/ml and 1.04 ug/ml, respectively, which increased to 0.733 ug/ml and 2.36 ug/ml, respectively when administered along with HREZ. Similarly, C_max_ and AUC_last_ for H/R/E/Z administered with meclizine were 5.25 ug/ml and 13.8 ug/ml*h for H, 8.8 ug/ml and 130.18 ug/ml*h for R, 1.92 ug/ml and 9.69 ug/ml*h for E and 63.1 ug/ml and 216.99 ug/ml*h for Z, respectively (Fig 7B-G). The plasma PK profiles of HREZ remained unchanged in the presence of meclizine. Comparative ratios of C_max_ and AUC_last_ for H, R, and Z with and without meclizine were close to 1 except for E, which showed minor interaction (C_max_ ratio: 1.41 and AUC_last_: 1.22) (Fig 8G). The average concentration of drug permeation in the lungs was determined at 6 and 24 h post-treatment, and the lung accumulation of HREZ in the presence or absence of meclizine remained comparable (Fig 8H). Similarly, HREZ did not affect lung deposition of meclizine at 6 and 24 h post-treatment (Fig 8H). Overall, the PK results suggested no adverse interactions between the HREZ plus meclizine combination vs. HREZ alone. Taken together, our study demonstrates an effect of meclizine on drug tolerance, no significant drug-drug interaction with HREZ, and a potentiating effect of meclizine on isoniazid. With over 50 years of safe clinical use of meclizine for nausea and vertigo [54, 62, 63], our findings suggest repurposing meclizine for developing a new combination for shortening therapy time for TB.

**Fig. 7.**
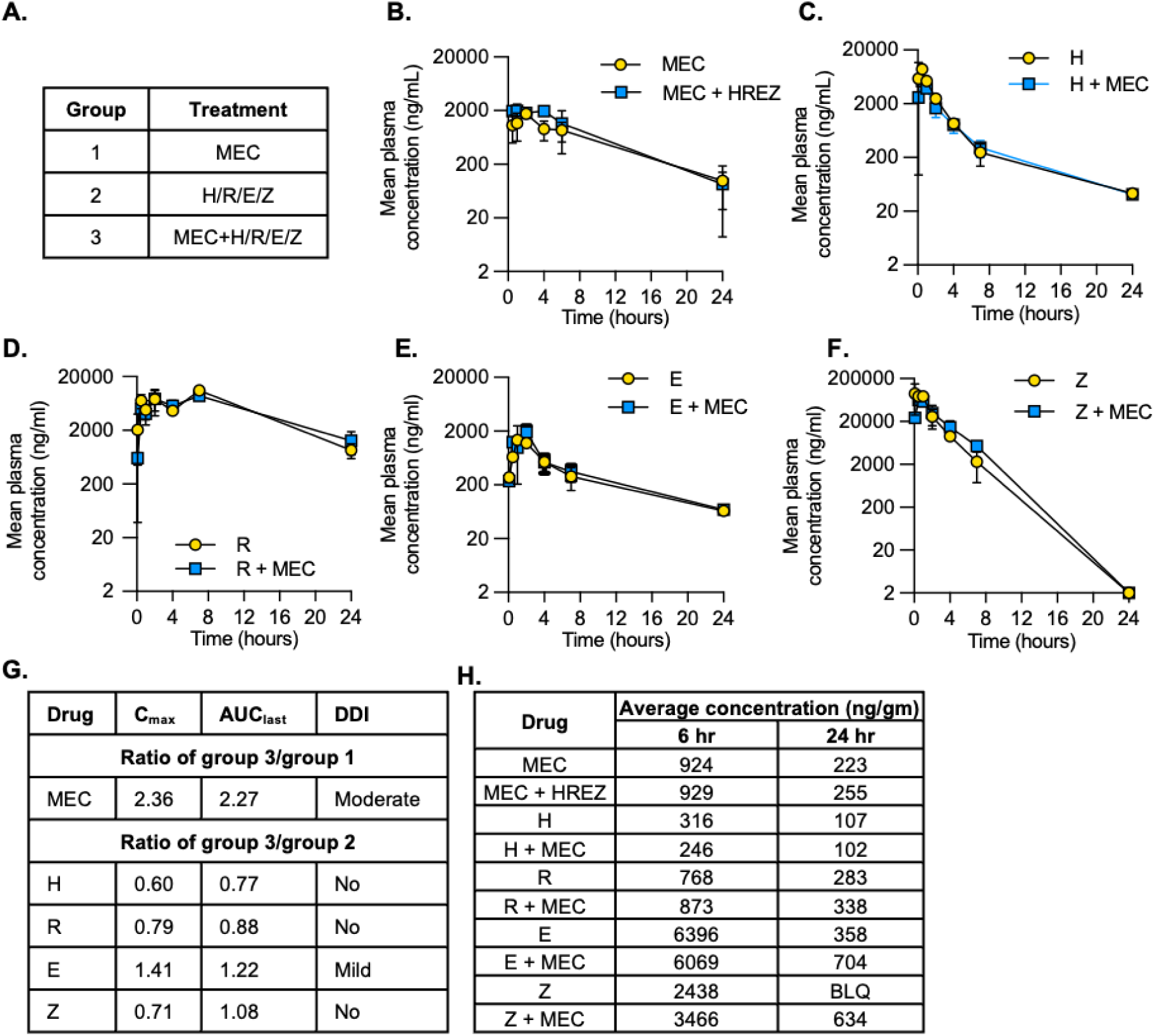
Meclizine exhibits no adverse interaction with anti-TB drugs. **A.** Three groups of treatment in BALB/c mice used in the pharmacokinetic study: MEC alone, front-line anti-TB combination therapy (HREZ), and combination (MEC + HREZ); (**B- F**) Line plots indicate pharmacokinetic profiles of MEC and individual drugs of the anti- TB therapy regimen analysed individually and in the presence of each other in the plasma of animals. Differences were non-significant by the Mann-Whitney test (p > 0.05). **G.** Ratios of C_max_ and AUC_last_ of individual drugs or a combination with meclizine to analyse drug-drug interactions. Doses used are the following: MEC, 25 mg/kg body weight, i.p.; H, 25 mg/kg body weight, p.o.; R, 10 mg/kg body weight, p.o.; E, 200 mg/kg body weight, p.o.; Z, 150 mg/kg body weight, orally; BDL, below the detection limit. All data are means ± SD of concentrations at each time point of samples in triplicates (n = 3 animals per group), **H.** Lung deposition of MEC alone and combined with anti-TB drugs at 6 h and 24 h post-intra-peritoneal administration.

## Discussion

Drug tolerance is commonly associated with diminished *Mtb* metabolism in response to unfavourable environmental conditions [65]. Consistent with this, studies have shown that IFNγ-mediated activation of macrophages *in vitro* and nitrosative stress encountered in AM/IM subsets in a murine infection model *in vivo* result in metabolic quiescence and drug tolerance in *Mtb* [4, 7]. However, human studies provide conclusive evidence that drug tolerance can be associated with metabolically active growing populations of *Mtb* [19, 20, 22, 66–69]. Maintenance of a metabolically active state confers an advantage to the pathogen during periods of relapse once antibiotic pressure is released [70, 71]. Consistent with this idea, we and others have demonstrated the emergence of drug tolerance in actively multiplying *Mtb* in naïve macrophages [20, 21]. This form of intra-macrophage drug tolerance relies on bacterial pathways that require active metabolism, such as efflux pumps, sulfur metabolism, and redox homeostasis (*e.g.,* trans-sulfuration pathway, Fe-S cluster biogenesis, and mycothiol biosynthesis) [20, 21, 72]. However, the aspects of macrophage physiology that engage with the pathogen to power bacterial sulfur and redox metabolism for drug tolerance remain poorly characterized.

In the work described above, we exploited a fluorescent *Mtb* redox reporter strain to functionally disentangle the relationship between macrophage bioenergetics and redox-mediated drug tolerance in actively multiplying *Mtb*. The redox reporter *Mtb* strain, which we have characterized previously [9, 21, 23, 73], demonstrates functional heterogeneity in BMDMs during the early stages of infection with respect to their transcriptional responses, energy metabolism, and mitoROS. BMDMs fraction relying on mitochondrial OXHPOS correlates with reductive shift and drug tolerance in *Mtb*, whereas the glycolytically active fraction promotes oxidative stress and drug lethality for *Mtb*. The significance of these two interconnected metabolic programs (OXPHOS vs. glycolysis) was demonstrated experimentally through genetic, biochemical, and pharmacological manipulation of macrophage bioenergetics that reduced redox heterogeneity and drug tolerance in *Mtb*.

Macrophage metabolism during *Mtb* infections displays complex patterns, as reports show a range of phenotypes, including balanced glycolysis and OXPHOS [74], elevated glycolysis [75, 76], and dampened glycolysis and OXPHOS [37, 77]. Our data added a new dimension to these studies by demonstrating the presence of macrophage subsets having relative variations in glycolysis vs. OXPHOS, which could be one of the explanations for inconsistencies in earlier studies. To our knowledge, the transcriptional and metabolic response (glycolytic and OXPHOS) have only been characterized in bulk murine BMDMs or human MDMs after infection with *Mtb* [37, 77], obscuring the response due to heterogeneity within the BMDM population. We confirmed the recent claim that *Mtb* infection decreases mitochondrial OXPHOS in bulk BMDMs compared to mock infection (Fig. S13). However, we phenotypically typed the infected BMDMs using the redox reporter *Mtb* strain before flow sorting, RNA-seq, and XF-flux analysis. This investigational and methodical pipeline revealed that infected BMDMs are composed of underlying fractions displaying differences in glycolysis, mitoROS, and OXPHOS.

In terms of metabolic parameters, we demonstrated that in BMDMs harboring *E*_MSH_- reduced bacteria, glucose is not preferentially converted to lactate as in BMDMs containing *E*_MSH_-oxidized bacteria or bystanders, but rather it is used to feed the OXPHOS pathway. Blocking glucose-linked OXPHOS by impairing the transport of pyruvate from cytosol to mitochondria induces oxidative stress and drug lethality in *Mtb*, whereas fuelling mitochondrial OCR by diverting glycolytic flux to OXPHOS maximizes a reductive shift and drug tolerance in *Mtb*. Similar to our findings, M1 and M2 activation programs are linked by divergent glucose flux into aerobic glycolysis and mitochondrial respiration, respectively [78]. Within the framework of *Mtb* infection, studies have reported that *Mtb* induces glycolysis in the lungs of infected mice [79] and lung granulomas from patients with active TB [80]. Interestingly, IFNγ-dependent control of *Mtb* relies on the induction of glycolytic enzymes by HIF1a in macrophages [81]. Despite the known role of glycolysis in restricting *Mtb* in these studies, the pathogen manages to shift the environment in its favor for persistence and drug tolerance during infection. Our findings raise the possibility that metabolic heterogeneity within macrophage populations regulates *Mtb’s* redox metabolism for survival and drug tolerance. Local factors found in lung macrophages may coordinate the metabolic network of macrophages during infection. In this context, a study using TB-pleural effusion (TB-PE) as a physiologically relevant fluid found in the human respiratory cavity showed that TB-PE *ex vivo* shifts the glycolysis-based metabolic program of macrophages to OXPHOS [82]. Given that a similar metabolic phenotype is exhibited by macrophages directly isolated from the pleural cavity of TB patients and lung biopsies of TB-infected non-human primates [83], we hypothesize that host energy metabolism and its correlation with redox-linked changes in sensitivity to anti-TB drugs, revealed by our study, is likely to be relevant in clinical settings of human TB.

Similar to *Mtb*, ROS/RNS and availability of nutrients in macrophages have been implicated in drug tolerance displayed by *S. aureus* and *Salmonella* [65, 71, 84–86]. Inflammasome-mediated activation of glycolysis in *S. aureus-*infected macrophages directly affected the metabolic state and the bacterium’s ability to enter into and exit from a drug-tolerant state [84]. Whether changes in the metabolic state of macrophages affect drug tolerance by impacting the redox metabolism of *S. aureus* and *Salmonella* remains to be characterized.

Why does *Mtb* experience oxidative stress in glycolytically active BMDMs? We found that in these BMDMs, mitoROS is enhanced, which serves as a signal to mobilize NADPH oxidase recruitment on mycobacterial phagosomes and promote xenophagy [38]. Consistent with this idea, elevated mitoROS in glycolytically active BMDMs correlated with higher cellular ROS. All these factors (mitoROS, NADPH oxidase, and xenophagy) could contribute to the oxidative shift in *Mtb* [23]. Diminished TCA cycle and ETC gene expression in glycolytically active BMDMs could result in reduced NADH production and electron shuttling through FET, resulting in elevated RET-ROS and oxidative shift in *Mtb*. Toll-like receptor (TLR) signaling alters respiratory complexes and generates RET-ROS at complex I [87]. Agreeing with this, genes associated with TLR signaling (e.g. TLR5, TLR9, TLR3, TLR1, MyD88) were marginally higher in glycolytically active BMDMs than in BMDMs having elevated OXPHOS (Table S1).

Our data also showed differential expression of the Hippo signaling pathway in flow-sorted BMDMs. Hippo-signaling promotes phagosomal and mitochondrial ROS production by enhancing the juxtaposition of mitochondria and phagosomes during bacterial infection [88]. Along with ROS generation, Hippo signaling maintains redox homeostasis in macrophages. A recent report demonstrated that Hippo kinases (MST1/2) function as ROS sensors and stabilize Nrf2 to attenuate overwhelming ROS generation [89]. The Mst1 receptor and Nrf2 pathway were induced in BMDMs that exhibited low mitoROS and harbored *Mtb* in a drug-tolerant *E*_MSH_-reduced state. Diminished expression of Nrf2 reversed these phenotypes, suggesting that MST-Nrf2 signaling is likely responsible for suppressing ROS generation in the host, thereby inducing a reductive shift and drug tolerance in *Mtb*. Increased expression of an Nrf2 signature in BMDMs containing drug-tolerant replicating bacteria overlapped with the reported expression of Nrf2 regulon in *Mtb* infected AM within 24 h of infection in mice [25] and in humans suffering from active TB [90]. Early expression of Nrf2 decreased pro-inflammatory pathways [25] and inhibited nitric oxide production [91, 92], which could allow *Mtb* to adopt a reductive redox state and proliferate in a permissive environment. We have previously shown that inhibition of iNOS induces a reductive shift in *E*_MSH_ of *Mtb* by reducing oxidative stress in *Mtb* [23].

Additionally, Nrf2-dependent induction of cystine-glutamate antiporter (xCT) could facilitate extracellular transport of cystine [93], thereby increasing intracellular cysteine, promoting a reductive shift in *E*_MSH_ of *Mtb* by utilization of host-derived cysteine into mycothiol. Since cysteine potentiates the activity of anti-TB drugs [73, 94], *Mtb* mitigates cysteine stress by increasing its utilization for the biogenesis of Fe- S clusters, H_2_S, and methionine metabolites in *E*_MSH_-reduced bacteria [21]. Our data and the expression of Nrf2 during mouse infection and in patients with active TB [25, 90, 95] imply that a similar correlation between Nrf2 signaling, host bioenergetics, and redox-linked drug tolerance in *Mtb* would emerge in human TB infection.

Our study presents opportunities to target redox-mediated drug tolerance in *Mtb*. Interfering with macrophage anti-oxidant response may facilitate improved bacterial killing by antibiotics. For example, nanoparticles containing Nrf2 activators sensitize intracellular killing of *S. aureus* in cultured macrophages [96]. A similar strategy could be used with Nrf2 inhibitors in potentiating the activity of anti-TB drugs. Although host redox balance is also important in cell-cell signaling, modulating anti-oxidant machinery may have harmful consequences unrelated to macrophages and the innate immune response [97, 98]. Our observation that shifting the metabolic reliance of macrophages from OXPHOS to glycolysis sensitizes *Mtb* to anti-TB drugs in macrophages opens the possibility of assessing agents that safely induce shifts in cellular metabolism during infection with *Mtb*. We focussed on an FDA-approved drug, meclizine, that suppresses OXPHOS by inhibiting the Kennedy pathway, leading to the accumulation of phosphoethanolamine that acts as a direct inhibitor of mitochondrial respiration [55]. Notably, suppression of OXPHOS resulted in RET-ROS and redirection of energy metabolism, leading to an oxidative shift in *E*_MSH_ of *Mtb* and elevated killing by anti-TB drugs. While the therapeutic potential of agents that target metabolic shifts in infected host cells is under-appreciated in bacterial infections, such strategies are favored in cancers [99], myocardial infarction [100], and stroke [101]. In this context, a recent adjunctive human clinical trial using an antidiabetic drug

(metformin) in combination with anti-TB drugs showed encouraging results in reducing inflammation and lung tissue damage [102]. Meclizine readily crosses the blood-brain barrier [103, 104], highlighting the potential therapeutic utility for treating TB meningitis. Meclizine has been used safely for >50 years for the treatment of nausea and vertigo [63, 64]. We found no adverse interaction of meclizine with first line anti-TB drugs in mice. Whether currently approved doses of meclizine achieve the required concentration in plasma and lung required for safe adjunctive effects with anti-TB drugs needs further experimentation. Safety studies are important because meclizine has a variable impact on OXPHOS in different cell types [54], and the associated increase in RET-ROS has been linked to necrosis of *Mtb*-infected macrophages [40]. Studies in animals, including non-human primates, have shown that higher doses of meclizine can be tolerated [105, 106]. Nonetheless, preclinical studies of efficacy and toxicity are required to determine optimal dosing and safety regimens before evaluating the utility of meclizine as an adjunct to anti-TB drugs in humans.

## Concluding Remarks

While targeting non-growing *Mtb* within the caseous centres of granuloma is being actively pursued in the TB field [107–111], the heterogeneity of the host macrophages (AM, IM, and BMDMs) and its impact on bacterial physiology and drug tolerance is only recently been getting attention. Adjuvant strategies using host-directed therapies to potentiate the efficacy of anti-TB drugs through targeting bioenergetic shifts in macrophages, such as those reported for meclizine, represent an important approach for future studies.

## Materials and Methods

### *Mtb* culture conditions

H37Rv strain of *Mtb* and H37Rv expressing Mrx1-roGFP2 (*Mtb*-roGFP2) were used in the study. The bacteria were grown to log phase in Middlebrook 7H9 broth supplemented with 10% albumin/dextrose/saline (ADS) or 10% oleic acid-albumin-dextrose catalase (OADC) supplement, 0.2% glycerol, 0.05% Tween 80, and 50μg/ml hygromycin B (only for *Mtb*-roGFP2).

### Bone Marrow-derived macrophages (BMDMs) culture and in vitro assays

BMDMs were isolated from C57BL/6J mice and cultured in DMEM (Cell Clone) supplemented with 10% FBS (Cell Clone), 30 ng/ml macrophage colony-stimulating factor (M-CSF, BioLegend), 2 mM L-glutamine (Sigma Aldrich), 1 mM sodium pyruvate (Sigma Aldrich), 10mM HEPES buffer (Sigma Aldrich), and 1% penicillin/streptomycin (Sigma Aldrich) at 37°C for 6 d. BMDM purity was checked by staining with monoclonal antibodies against MerTK (Thermo Fischer, (DS5MMER) Alexa Fluor 700) and CD64(anti-mouse, BioLegend). For FACS-based experiments, BMDMs were infected with *Mtb*-roGFP2 at multiplicity of infection (moi) of 10 for 3h, before extracellular bacteria were removed by washing thrice with Dulbecco’s phosphate-buffered saline (DPBS). Fresh optiMEM medium (Thermo-Fischer) supplemented with 5% FBS and 30ng/ml M-CSF was added, and cells were incubated for different time points according to experimental requirements.

For CFU determination, cells were infected at a moi of 2 with *Mtb* H37Rv for 3h and lysed with 0.05% SDS in water and plated on 7H11 agar supplemented with 10% OADC at different time points post-infection. Cells were incubated in optiMEM medium supplemented with 5% FBS and 30ng/ml M-CSF post-infection, and fresh medium was added every 48 hours.

### Redox profiling of intracellular *Mtb*-roGFP2

At different time points post-infection, cells were washed and scraped off in 1X DPBS and acquired using BD FACS Aria Fusion flow cytometer and data was analysed using BD FACS Diva software. At least 10,000 GFP-positive events were analysed by excitation at 405 and 488 nm with a constant emission (510 nm) to determine *E*_MSH_ under different treatment conditions. Events were gated based on a previously developed strategy [23], where infected BMDMs were treated with 10 mM CHP for complete oxidation or 100 mM DTT for complete reduction of the biosensor.

### Flow sorting of macrophages infected with *Mtb*-roGFP2

Post-infection at the required time points, the growth medium was removed from the cells, followed by washing with DPBS supplemented with 2% FBS. Fresh DPBS was added, and the cells were gently scrapped off using a cell scrapper. The cell suspension was centrifuged, and cells were resuspended in DPBS and passed through a 40-micron (40μ) cell strainer. Cells were sorted using BD FACS Aria flow cytometer (BD Biosciences) employing the 405 nm and the 488 nm lasers into *E*_MSH_- reduced, -basal, -oxidized and bystander sub-populations. Cells were gated using 10 mM CHP (oxidant) and 100 mM DTT (reductant) into *E*_MSH_-oxidized (∼-240 mV) and - reduced (∼-300 mV), respectively. The GFP-negative population was gated as bystanders. The sorting was done at a “four-way purity” setting, and the purity of sorting was assessed by post-sort analysis. The uninfected cells were also treated with the same conditions as the infected and mock-sorted to eliminate differences that may arise due to cell sorting.

### RNA isolation, library preparation for RNA-Seq

RNA sequencing was done at the Next-Gen Genomics Facility at the National Centre for Biological Sciences (NCBS), Bangalore. Total RNA was extracted from FACS-sorted macrophage sub-populations using Qiagen RNeasy Mini Kit. All RNA samples (three biological replicates in each group) passed quality control analysis on Agilent 4200 TapeStation system. rRNA depletion was done using NEBNext® rRNA Depletion Kit (Human/Mouse/Rat) with RNA Sample Purification Beads (E6350X). Directional poly-A+ RNA-sequencing libraries were generated with the NEBNext® Ultra™ II Directional RNA Library Prep with Sample Purification Beads (E7765L). Libraries were sequenced on a NovaSEQ6000 (Illumina) using a paired end sequencing format with read length of 100 bp.

### RNA-Seq Data Analysis for macrophage subpopulations

RNA-Seq data was preprocessed to obtain read counts using the “htseq” pipeline in Python [112] using the reference genome of *Mus musculus* (GCF_000001635.27_GRCm39). Differential gene expression analysis was performed on the different subgroups with a base mean value of expression greater than 1, log2 fold change greater than 0.6 and an adjusted p-value less than 0.1 using the DESeq2 pipeline [113]. To characterize the differentially expressed genes between the oxidized and reduced populations, DESeq2 was used to identify genes that were specifically upregulated in each category with respect to the other category using the above criteria. Similarly, gene signatures pertaining to bystander, reduced and oxidized populations with respect to the reference uninfected genes were also calculated using the above criteria. Gene expression values for genes common with differentially expressed genes and different gene signatures (see gene lists section) were plotted as heatmaps using cluster map function in the seaborn package after conversion of read counts to transcripts per million (TPM) followed by z-scoring across the experimental conditions to avoid biases due to absolute values of gene expression.

Transcriptomic data are uploaded to the NCBI GEO database, accession number GSE283664.

### Gene lists and overlap analysis

Gene list belonging to the oxidative phosphorylation pathway was obtained from the MSigDB database [114]. Gene list belonging to the antioxidant pathway was obtained from Gene Ontology Molecular Functions subcategory. The Hippo signaling pathway was obtained from Reactome gene sets through the MSigDB database[114]. M1 and M2 specific macrophage polarization signatures were obtained from GSE5099 [115] through the MSigDB database. Gene signatures characterizing the transcriptomic states post 4 hours and 24 hours were obtained from [24]. AM and IM-specific gene signatures upon infection with Mycobacterium tuberculosis were obtained from [25] (greater than log2 fold change). Finally, gene signatures belonging to the Nrf2 pathway were obtained from [25]. Gene lists corresponding to Nrf2 chip-seq performed on peritoneal macrophages with differential peaks and associated Nrf2 motifs were obtained from GSE75177 [25, 43]. Overlap analysis between the different gene signatures and pathways were performed using the *ggVennDiagram* module in R. Quantification of pathway activity followed by z-normalization was performed using the ssGSEA function in *gseapy* library in Python [116].

### MIC and MBC determination

*Mtb* H37Rv were cultured in 7H9+ADS medium till mid-log phase (∼0.4-0.6 OD_600_) post which ∼10^6^ *Mtb* were seeded in 200 ul 7H9+ADS medium without Tween-80. After 5 days of incubation at 37°C, 20 μL of Alamar blue was added, and the plates were re-incubated for 24 h. The fluorescence readings were recorded in a Spectramax M3 microplate reader with excitation at 530 nm and emission at 590 nm. Percentage inhibition was calculated based on the relative fluorescence units and the minimum concentration that resulted in at least 90% inhibition (Minimum Inhibitory Concentration or MIC).

For determining the minimum bactericidal concentration (MBC), 20μl of cells from the MIC plate were spotted on 7H11-agar plates supplemented with 10% OADC. The plates were incubated at 37°C for 3 weeks and MBC was defined as the concentration where no bacterial colonies were visible on the plates.

### ROS measurement assays

For ROS measurement assays, BMDMs were seeded in 24-well plates (0.3 million cells/well). To measure cellular ROS, cells were stained with 2.5μM CellROX Deep Red dye (Thermo Fischer Scientific-Life technologies) for 30 minutes in Dulbecco’s phosphate-buffered saline (DPBS). As positive control, cells were pretreated with 100μM Menadione (Sigma Aldrich) for 1 hour before staining. For mitochondrial ROS measurement, cells were stained with 2.5 μM MitoSOX Red dye (Thermo Fischer Scientific-Life technologies) for 30 mins in the HBSS buffer. As a positive control for mitochondrial ROS measurement, cells were treated with 20μM Antimycin A (Sigma Aldrich) for the last 15 minutes of staining. After completion of staining, cells were washed thrice with DPBS, and fluorescence was assessed using BD FACS Aria flow cytometer (BD Biosciences) at prescribed excitation and emission maxima CellROX DeepRed: 644/665; MitoSOX Red: 544/610.

### siRNA-mediated knockdown of Nrf2 in BMDMs

BMDMs were seeded in a 6-well plate after 6 days of differentiation. Post reattachment, the medium was replaced with OptiMEM with 40nM Nrf2-targeting siRNA (siNRF2) or scrambled siRNA negative control (siSCR) (Integrated DNA Technologies) and 4ul Lipofectamine 3000 per well (Thermo-Fischer Scientific). The sequence of siRNA against Nrf2 was:

*Sequence: rGrCrG rArUrG rArArU rUrUrU rArUrU rCrUrG rCrUrU rUrCA T*

*Sequence: rArUrG rArArA rGrCrA rGrArA rUrArA rArArU rUrCrA rUrCrG rCrCrA*

24 hours post-transfection, cells were infected with *Mtb-*roGFP2 or *Mtb* H37Rv according to experimental requirements and downstream assays were performed.

### qRT-PCR

Infected BMDMs were lysed in the RLT buffer (Qiagen), and total RNA was isolated according to the manufacturer’s protocol using the Qiagen RNeasy Plus Mini Kit. Isolated RNA was further treated with DNase I (Thermo Scientific) to remove any residual genomic DNA. 500ng of RNA was used to synthesize cDNA for downstream qRT-PCR analysis using the iScript Select cDNA Synthesis Kit (BioRad). (CFX96 RT-PCR System, BioRad) and iQ SYBR Green Supermix (BioRad) were used to assess gene expression, and the primers used for qRT-PCR for different genes were as follows:

Nrf2 FP 5’-CAGCATAGAGCAGGACATGGAG-3’

Nrf2 RP 5’-GAACAGCGGTAGTATCAGCCAG-3’

Gstm1 FP 5’-TGTTTGAGCCCAAGTGCCTGGA-3’

Gstm1 RP 5’-TAGGTGTTGCGATGTAGCGGCT-3’

GclcFP 5’-ACACCTGGATGATGCCAACGAG-3’

Gclc RP 5’-CCTCCATTGGTCGGAACTCTAC-3’

Pik3cb FP 5’-CAGTTTGGTGTCATCCTGGAAGC-3’

Pik3cb RP 5’-TCTGCTCAGCTTCACCGCATTC-3’

Gapdh FP 5’-CATCACTGCCACCCAGAAGACTG-3’

Gapdh RP 5’-ATGCCAGTGAGCTTCCCGTTCAG-3’

Mouse Gapdh expression (Ct values) was used as the normalization control in all cases.

### Antibiotic tolerance assay

BMDMs were infected at a moi of 2 with *Mtb* H37Rv for 3h. Extracellular bacteria was removed by washing the cells thrice with DPBS, and treatment was initiated (SFN, MEC, glucose/galactose, UK5099). At the required time points, cells were lysed with 0.05% SDS and the lysate was plated on 7H11 agar plates supplemented with OADC after serially diluting the lysate. The plates were incubated at 37°C and colony forming units were enumerated after 3 weeks.

### MTT assay

BMDMs were seeded in a 96-well plate (30000 cells/ well), and upon attachment to the surface, they were treated with different concentrations of drugs for 24 h. post- treatment, 0.25 mg/ml (3-(4,5-dimethylthiazol-2-yl)-2,5-diphenyltetrazolium bromide (MTT) was added to the cells and incubated for 2.5 hours at 37°C and 5% CO_2_. To assess survival, absorbance was measured at 560nm using a Spectramax plate reader. Untreated cells or cells treated with 50% DMSO for 10 minutes were taken as positive and negative controls, respectively.

### Oxygen consumption rate (OCR) measurement/Modified Mitostress test

Before the assay, the Seahorse XFp Cell Culture plate was coated with 15 μl Cell-Tak reagent (5 μl Cell-Tak, 145 μl 0.1N sodium bicarbonate at pH 8) for 30 minutes at 37°C. After coating, BMDMs were seeded at a density of 40000 cells per well in the cell culture plate in Agilent Seahorse Mitostress test medium supplemented with 2mM L-Glutamine, 10mM D-Glucose and 1mM Sodium Pyruvate.

Three OCR readings were taken under basal conditions followed by periodic addition of oligomycin (2 μM) and a combination of rotenone (2 μM) and antimycin (4 μM). Measurements were taken using an 8-well Seahorse XFp analyzer (Agilent Technologies). The data was analysed using GraphPad Prism ver. 10. Basal respiration was calculated by subtracting the non-mitochondrial OCR (lowest reading after adding rotenone and antimycin) from the last reading before oligomycin injection. ATP production was calculated by subtracting the lowest reading after oligomycin injection from the previous reading before oligomycin injection. Proton leak was calculated by subtracting the non-mitochondrial OCR from the lowest reading after oligomycin injection.

To assess respiratory parameters in the presence of 10 mM glucose or galactose, cells were incubated with either sugar source in the medium for 24 h, after which the modified mitostress test was performed. For different inhibitors/drugs, treatment was done in regular culture plates post, and the cells were seeded in the Seahorse cell culture plates for infection.

### Extracellular Acidification Rate (ECAR) measurement

BMDMs were seeded in a Cell-Tak coated Seahorse XFp Cell Culture plate (40000 cells per well) in Agilent Mitostress test medium supplemented with 2mM L-Glutamine. The standard Agilent Seahorse Glycolytic Stress test was run according to the manufacturer’s protocol wherein three ECAR readings each were measured upon sequential addition of 10 mM D-Glucose, 2 μM oligomycin and 100 mM 2- deoxyglucose (2-DG). For different drugs, macrophages were seeded directly into the XFp plates, and the glycolysis stress test was run. Glycolysis was calculated by subtracting the lowest reading before glucose injection (non-glycolytic acidification) from the highest reading post glucose injection. Glycolytic capacity was calculated by subtracting the lowest reading before glucose injection from the highest reading after oligomycin injection.

### Mitochondrial membrane polarization

∼0.3 million BMDMs in a 24-well plate were infected with *Mtb* H37Rv, and at 24 h post-infection, cells were washed with DPBS and stained with 2 μM JC-1 dye (Thermo Scientific). After 30 minutes of incubation at 37°C, cells were washed three times with DPBS, and fluorescence was measured using BD FACS Aria Fusion flow cytometer at 485/535 nm and 550/600 nm excitation/emission wavelengths. Cells were treated with 50 μM FCCP for 30 minutes during staining as positive control.

### Mitochondrial imaging

For imaging of murine BMDMs, bone marrow was obtained from the femurs of 6- to 8- 8-week-old female C57BL/6 mice (Charles River). Animal protocols were reviewed and approved by EPFL’s Chief Veterinarian, by the Service de la Consommation et des Affaires Vétérinaires of the Canton of Vaud, and by the Swiss Office Vétérinaire Fédéral under license VD 3434. Bone marrow was cultured in Dulbecco’s Modified Eagle Medium (DMEM) (Gibco) supplemented with 10% FBS and 20% L929-cell-conditioned medium (as a source for macrophage-colony stimulating factor protein or M-CSF) for 7 days. No antibiotics were used in the cell culture media for all cell types to avoid activation of macrophages or inhibition of *Mtb* growth.

Differentiated BMDMs were seeded in 24 well plates (Ibidi) suitable for high-resolution imaging. A 1 mL aliquot of a culture of the *Mtb* H37Rv mCherry strain grown to exponential phase (OD_600_ 0.3-0.5) was centrifuged at 5000 g for 5 minutes at room temperature, the supernatant was removed, and the cell pellet was resuspended in the macrophage media. A single cell suspension was generated via filtration through a 5-μm syringe filter (Millipore). BMDMs were infected with a 200-fold dilution of the single-cell suspension for 4 hours. Infected cells were washed with pre-warmed PBS to remove extracellular bacteria and incubated in the presence or absence of 20 μM meclizine for 24 hours at 37°C in 5% CO_2_.

For staining of mitochondria, cells were washed with a pre-warmed medium and incubated with 50 nM MitoTracker Deep Red FM (Invitrogen, M22426) for 1 hour. Stained cells were then washed to remove excess dye and fixed with freshly prepared 4% paraformaldehyde at room temperature for a minimum of 2 hours. Fixed cells were washed to remove excess fixative and stained with Hoechst (Thermo Fisher) at 5 μg/mL for 30 minutes at room temperature for visualizing nuclei.

Confocal images were acquired on a Leica SP8 inverted microscope using a 63x oil objective (NA=1.4). Z stacks were subsequently deconvolved using the Huygens Professional Deconvolution software (Scientific Volume Imaging). ImageJ and Imaris 10.1 (Bitplane) were used to quantify mitochondrial volume and surface area. Imaris was also used for rendering 3D images.

### RNA isolation, sequencing and data analysis for meclizine-treated macrophages

Differentiated BMDMs were infected with *Mtb*-roGFP2 at a moi 10 for 3 hours, after which the extracellular bacteria were removed by washing thrice with DPBS. Post-infection, cells were treated with 0.2% DMSO (solvent control) or 20 μM meclizine hydrochloride (Sigma-Aldrich). At 24 h p.i., infected BMDMs from each condition were flow-sorted using GFP as a marker for infection and total RNA was isolated and subjected to sequencing as explained earlier.

Raw reads were obtained as fastq files. The reference genome sequence and annotation files for mouse (*Mus musculus*, GCF_000001635.27_GRCm39) were downloaded from the NCBI ftp (“ftp.ncbi.nlm.nih.gov”). The raw read quality was checked using the FastQC software (version v0.11.5), and the differential gene expression (DGE) analysis was done using the EdgeR package in R [117]. Transcriptomic data are uploaded to the NCBI GEO database, accession number GSE283874.

### *In vivo* mice experiment

Six to seven weeks old C3HeB/FeJ mice were infected with *Mtb* H37Rv through the aerosol route using a Madison Chamber infection instrument with ∼30-40 CFUs. After 2 weeks of infection progression, mice were either treated with vehicle control (1XPBS + 5% ethanol + 5% Cremophor EL-25) or 25 mg/kg/d meclizine hydrochloride (Sigma-Aldrich). Post 4 weeks of infection, six mice were sacrificed to assess the CFU load in the lungs and 10 mg/kg/d isoniazid was introduced into the treatment regimen for the rest, dividing mice into four treatment groups- (i) vehicle treatment, (ii) isoniazid treatment (10 mg/kg/d), (iii) meclizine treatment (25 mg/kg/d), and (iv) meclizine plus isoniazid treatment. At 10 weeks post-infection, mice were euthanized, and lungs were harvested to analyse gross pathology, histopathology and CFU load. The upper right lobe of the lungs of animals from each group was fixed in 10% neutral-buffered formalin. Fixed tissues were prepared as 5-um-thick sections, embedded in paraffin, and stained with hematoxylin and eosin. Tissue sections were coded, and a certified pathologist analyzed coded sections to assess for granuloma formation and lung damage [21, 118, 119]. Remaining tissue samples were homogenized in 2 ml of sterile DPBS, serially diluted, and plated on 7H11-OADC agar plates supplemented with lyophilized BBL MGIT PANTA antibiotic mixture (polymyxin B, amphotericin B, nalidixic acid, trimethoprim, and azlocillin, as supplied by BD). Plates were incubated at 37°C for 3 weeks before colonies were enumerated.

### Lung tissue preparation and macrophage staining

At 10 weeks post-infection aseptically removed lungs of the mice from the four treatment groups minced and digested in serum-free DMEM containing 0.2 mg/mL Liberase DL (Roche) and 0.1 mg/mL DNase I (Roche) for 60 minutes at 37°C at 180 rpm. This was followed by mechanical disruption of the suspension using the GentleMACS dissociator (Militenyi Biotech) and de-clumping by passage through 70μ nylon cell strainers (Corning). To remove all red blood cells, the single cell suspension was incubated in RBC lysis buffer for 15 mins with gentle vortexing [21].

For staining of surface markers, lung cell suspensions were blocked with anti-mouse CD16/CD32 (Fc Block, BD Pharmingen) followed by anti-mouse CD64 (FCγRI, APC-conjugated; BioLegend), MerTK (DS5MMER, Alexa Fluor 700-conjugated; eBioscience) antibodies. The CD64 and MerTK double-positive cells were gated to analyse the total lung macrophage population in mice lungs (109). Alveolar and interstitial macrophages (AMs and IMs) were classified based on Siglec F (PerCP-conjugated; BioLegend).

### PK-PD DDI studies

4-6 weeks old BALB/c mice (n = 6 per group) were dosed with meclizine, first-line anti-TB therapy regimen HREZ or a combination of both, as follows: MEC, 25 mg/kg body weight intraperitoneal (ip) injection; INH (H) 25 mg/kg body weight orally; RIF (R) 10 mg/kg body weight orally; EMB (E) 200 mg/kg body weight orally and PZA (Z) 150 mg/kg body weight orally. HREZ were made in 0.5% hydroxypropyl methylcellulose and 0.1% Tween 80, and meclizine was prepared in 5% Cremophor EL 25 (Merck). Blood samples were collected from animals in the three treatment groups at regular intervals over 24 h from dosing, and plasma was isolated. All plasma samples, along with drug controls, were analysed by LC-MS (Waters Acquity UPLC system, flow rate 0.300 μl/mL with gradient elution; Waters Acquity-TQD Triple Quadrupole Mass Spectrometer). Calibration curves were generated for each analyte and accepted if 67% of calibration points were within ± 20% nominal concentration.

The pharmacokinetics profile was generated for each drug in terms of mean plasma concentration over 24 h, maximum concentration achieved (C_max_) and net exposure (AUC_last_) [120].

### Statistical analysis

Statistical tests and the generation of graphs were done in GraphPad Prism 10.2.0. A description of statistical tests can be found in the figure legends. Significance was defined as p<0.05.

### Ethics statement

This study was carried out strictly following the guidelines provided by the Committee for the Purpose of Control and Supervision on Experiments on Animals (CPCSEA), Government of India. The protocol for the animal experiment was approved by the Animal Ethical Committee on the Ethics of Animal Experiments, IISc, Bangalore, India (approval number: CAF/Ethics/972/2023). All humane efforts were made to minimize the suffering.

## Acknowledgements

We thank Prof. Karl Drlica at the Public Health Research Institute (PHRI), New Jersey Medical School, for the critical reading of the manuscript and his valuable input. We acknowledge Inder Raj Singh at NCBS, Bengaluru, for his assistance in analysing the RNA-Seq data. We also acknowledge the CIDR BSL-3 facility, Indian Institute of Science (IISc), for carrying out experiments with *Mtb* and Dr. Awadhesh Pandit and the NGS facility, NCBS, for RNA-sequencing. RM and VVT thank the BioImaging and Optics Platform Core Facility at EPFL and the Infectious Diseases Imaging Platform (IDIP) at the Centre for Integrated Infectious Diseases at the Heidelberg University Medical Faculty. VVT acknowledges support from the Holcim Stiftung zur Förderung der Wissenschaftlichen and Heidelberg University Medical Faculty.

## Author contributions

Conceptualization: VY and AS. Methodology: VY, RM, RSR, SSM. Bioinformatic analysis: VY, SS, NM, MJ, ASNS, and AS. Investigation: VY, RM, RSR, SSM. Supervision: VY and AS. Data analysis: VY, SS, NM, RM, RSR, SSM, RKS, SN, VVT, MJ, ASNS, AS. Funding acquisition: AS. Writing-original draft: VY and AS. Writing- review and editing: VY and AS. All authors read and approved the final manuscript.

## Data availability

The RNA-sequencing data presented in the manuscript are attached in the supplementary tables (Table S1-S7) and submitted to the NCBI Gene Expression Omnibus (GEO). Reviewers can access the data files using the tokens: Redox subpopulations- GSE283664: ujsnwgwkrxaznsz and meclizine treatment- GSE283874: wzwhymicrhglvyl

## References

1. Lenaerts, A., C.E. Barry, 3rd, and V. Dartois, *Heterogeneity in tuberculosis pathology, microenvironments and therapeutic responses*. Immunol Rev, 2015. 264(1): p. 288–307.

2. Mishra, R., et al., Heterogeneous Host-Pathogen Encounters Coordinate Antibiotic Resilience in Mycobacterium tuberculosis. Trends Microbiol, 2021. 29(7): p. 606–620.

3. Cadena, A.M., S.M. Fortune, and J.L. Flynn, Heterogeneity in tuberculosis. Nat Rev Immunol, 2017. 17(11): p. 691–702.

4. Liu, Y., et al., Immune activation of the host cell induces drug tolerance in Mycobacterium tuberculosis both in vitro and in vivo. J Exp Med, 2016. 213(5): p. 809–25.

5. Huang, L., et al., Growth of Mycobacterium tuberculosis in vivo segregates with host macrophage metabolism and ontogeny. J Exp Med, 2018. 215(4): p. 1135–1152.

6. Pisu, D., et al., Dual RNA-Seq of Mtb-Infected Macrophages In Vivo Reveals Ontologically Distinct Host-Pathogen Interactions. Cell Rep, 2020. 30(2): p. 335–350 e4.

7. Pisu, D., et al., Single cell analysis of M. tuberculosis phenotype and macrophage lineages in the infected lung. J Exp Med, 2021. 218(9).

8. Kalam, H., et al., Identification of host regulators of Mycobacterium tuberculosis phenotypes uncovers a role for the MMGT1-GPR156 lipid droplet axis in persistence. Cell Host Microbe, 2023. 31(6): p. 978–992 e5.

9. Das, M., et al., Cysteine desulfurase (IscS)-mediated fine-tuning of bioenergetics and SUF expression prevents Mycobacterium tuberculosis hypervirulence. Sci Adv, 2023. 9(50): p. eadh2858.

10. Anand, K., et al., Mycobacterium tuberculosis SufR responds to nitric oxide via its 4Fe-4S cluster and regulates Fe-S cluster biogenesis for persistence in mice. Redox Biol, 2021. 46: p. 102062.

11. Rutschmann, O., C. Toniolo, and J.D. McKinney, Preexisting Heterogeneity of Inducible Nitric Oxide Synthase Expression Drives Differential Growth of Mycobacterium tuberculosis in Macrophages. mBio, 2022. 13(5): p. e0225122.

12. Sarathy, J.P., et al., Extreme Drug Tolerance of Mycobacterium tuberculosis in Caseum. Antimicrob Agents Chemother, 2018. 62(2).

13. Hicks, N.D., et al., Clinically prevalent mutations in Mycobacterium tuberculosis alter propionate metabolism and mediate multidrug tolerance. Nat Microbiol, 2018. 3(9): p. 1032–1042.

14. Muñoz-Elías, E.J. and J.D. McKinney, Mycobacterium tuberculosis isocitrate lyases 1 and 2 are jointly required for in vivo growth and virulence. Nat Med, 2005. 11(6): p. 638–44.

15. Safi, H., et al., Phase variation in Mycobacterium tuberculosis glpK produces transiently heritable drug tolerance. Proceedings of the National Academy of Sciences, 2019. 116(39): p. 19665–19674.

16. Sharma, A., et al., VapC21 Toxin Contributes to Drug-Tolerance and Interacts With Non-cognate VapB32 Antitoxin in Mycobacterium tuberculosis. Frontiers in Microbiology, 2020. 11.

17. Singh, R., C.E. Barry, and H.I.M. Boshoff, The Three RelE Homologs of Mycobacterium tuberculosis Have Individual, Drug-Specific Effects on Bacterial Antibiotic Tolerance. Journal of Bacteriology, 2010. 192(5): p. 1279–1291.

18. Tiwari, P., et al., MazF ribonucleases promote Mycobacterium tuberculosis drug tolerance and virulence in guinea pigs. Nature Communications, 2015. 6(1): p. 6059.

19. Manina, G., N. Dhar, and J.D. McKinney, Stress and host immunity amplify Mycobacterium tuberculosis phenotypic heterogeneity and induce nongrowing metabolically active forms. Cell Host Microbe, 2015. 17(1): p. 32–46.

20. Adams, K.N., et al., Drug tolerance in replicating mycobacteria mediated by a macrophage-induced efflux mechanism. Cell, 2011. 145(1): p. 39–53.

21. Mishra, R., et al., Targeting redox heterogeneity to counteract drug tolerance in replicating Mycobacterium tuberculosis. Sci Transl Med, 2019. 11(518).

22. Ufimtseva, E.G. and N.I. Eremeeva, Drug-Tolerant Mycobacterium tuberculosis Adopt Different Survival Strategies in Alveolar Macrophages of Patients with Pulmonary Tuberculosis. Int J Mol Sci, 2023. 24(19).

23. Bhaskar, A., et al., Reengineering redox sensitive GFP to measure mycothiol redox potential of Mycobacterium tuberculosis during infection. PLoS Pathog, 2014. 10(1): p. e1003902.

24. Andreu, N., et al., Primary macrophages and J774 cells respond differently to infection with Mycobacterium tuberculosis. Scientific Reports, 2017. 7(1): p. 42225.

25. Rothchild, A.C., et al., Alveolar macrophages generate a noncanonical NRF2- driven transcriptional response to Mycobacterium tuberculosis in vivo. Sci Immunol, 2019. 4(37).

26. Ge, S.X., D. Jung, and R. Yao, ShinyGO: a graphical gene-set enrichment tool for animals and plants. Bioinformatics, 2020. 36(8): p. 2628–2629.

27. Kuleshov, M.V., et al., Enrichr: a comprehensive gene set enrichment analysis web server 2016 update. Nucleic Acids Res, 2016. 44(W1): p. W90–7.

28. Keenan, A.B., et al., ChEA3: transcription factor enrichment analysis by orthogonal omics integration. Nucleic Acids Research, 2019. 47(W1): p. W212–W224.

29. Waxman, E.A., Bach2 is a potent repressor of Nrf2-mediated antioxidant enzyme expression in dopaminergic neurons. bioRxiv, 2019: p. 687590.

30. Tsukumo, S., et al., Bach2 maintains T cells in a naive state by suppressing effector memory-related genes. Proc Natl Acad Sci U S A, 2013. 110(26): p. 10735–40.

31. Mo, J.S., et al., Cellular energy stress induces AMPK-mediated regulation of YAP and the Hippo pathway. Nat Cell Biol, 2015. 17(4): p. 500–10.

32. Wang, W., et al., AMPK modulates Hippo pathway activity to regulate energy homeostasis. Nature Cell Biology, 2015. 17(4): p. 490–499.

33. Biswal, P., et al., The interplay between hippo signaling and mitochondrial metabolism: Implications for cellular homeostasis and disease. Mitochondrion, 2024. 76: p. 101885.

34. Dinkova-Kostova, A.T. and A.Y. Abramov, The emerging role of Nrf2 in mitochondrial function. Free Radical Biology and Medicine, 2015. 88: p. 179–188.

35. Holmström, K.M., et al., Nrf2 impacts cellular bioenergetics by controlling substrate availability for mitochondrial respiration. Biology Open, 2013. 2(8): p. 761–770.

36. Kim, T.H., et al., NRF2 blockade suppresses colon tumor angiogenesis by inhibiting hypoxia-induced activation of HIF-1α. Cancer Res, 2011. 71(6): p. 2260–75.

37. Cumming, B.M., et al., Mycobacterium tuberculosis induces decelerated bioenergetic metabolism in human macrophages. Elife, 2018. 7.

38. Chandra, P., et al., Inhibition of Fatty Acid Oxidation Promotes Macrophage Control of Mycobacterium tuberculosis. mBio, 2020. 11(4).

39. Murphy, Michael P., How mitochondria produce reactive oxygen species. Biochemical Journal, 2008. 417(1): p. 1–13.

40. Roca, F.J., et al., Tumor necrosis factor induces pathogenic mitochondrial ROS in tuberculosis through reverse electron transport. Science, 2022. 376(6600): p. eabh2841.

41. Esteras, N. and A.Y. Abramov, Nrf2 as a regulator of mitochondrial function: Energy metabolism and beyond. Free Radical Biology and Medicine, 2022. 189: p. 136–153.

42. Ryan, D.G., et al., Nrf2 activation reprograms macrophage intermediary metabolism and suppresses the type I interferon response. iScience, 2022. 25(2): p. 103827.

43. Otsuki, A., et al., Unique cistrome defined as CsMBE is strictly required for Nrf2- sMaf heterodimer function in cytoprotection. Free Radic Biol Med, 2016. 91: p. 45–57.

44. Ren, X., et al., Overcoming the compensatory elevation of NRF2 renders hepatocellular carcinoma cells more vulnerable to disulfiram/copper-induced ferroptosis. Redox Biology, 2021. 46: p. 102122.

45. Li, Y., et al., Sorafenib induces mitochondrial dysfunction and exhibits synergistic effect with cysteine depletion by promoting HCC cells ferroptosis. Biochem Biophys Res Commun, 2021. 534: p. 877–884.

46. Dézsi, C.A., Trimetazidine in Practice: Review of the Clinical and Experimental Evidence. Am J Ther, 2016. 23(3): p. e871–9.

47. Shi, L., et al., Biphasic Dynamics of Macrophage Immunometabolism during Mycobacterium tuberculosis Infection. mBio, 2019. 10(2): p. 10.1128/mbio.02550-18.

48. Russell, D.G., et al., Foamy macrophages and the progression of the human tuberculosis granuloma. Nat Immunol, 2009. 10(9): p. 943–8.

49. Guerrini, V., et al., Storage lipid studies in tuberculosis reveal that foam cell biogenesis is disease-specific. PLoS Pathog, 2018. 14(8): p. e1007223.

50. McCommis, K.S. and B.N. Finck, Mitochondrial pyruvate transport: a historical perspective and future research directions. Biochem J, 2015. 466(3): p. 443–54.

51. Miyajima, H., T. Oda, and A. Ichiyama, Induction of mitochondrial serine:pyruvate aminotransferase of rat liver by glucagon and insulin through different mechanisms. J Biochem, 1989. 105(4): p. 500–4.

52. Martino, M.R., et al., Silencing alanine transaminase 2 in diabetic liver attenuates hyperglycemia by reducing gluconeogenesis from amino acids. Cell Reports, 2022. 39(4): p. 110733.

53. Höfler, S., et al., Dealing with the sulfur part of cysteine: four enzymatic steps degrade l-cysteine to pyruvate and thiosulfate in Arabidopsis mitochondria. Physiologia Plantarum, 2016. 157(3): p. 352–366.

54. Gohil, V.M., et al., Nutrient-sensitized screening for drugs that shift energy metabolism from mitochondrial respiration to glycolysis. Nat Biotechnol, 2010. 28(3): p. 249–55.

55. Gohil, V.M., et al., Meclizine inhibits mitochondrial respiration through direct targeting of cytosolic phosphoethanolamine metabolism. J Biol Chem, 2013. 288(49): p. 35387–95.

56. Wai, T. and T. Langer, Mitochondrial Dynamics and Metabolic Regulation. Trends in Endocrinology & Metabolism, 2016. 27(2): p. 105–117.

57. Liesa, M. and Orian S. Shirihai, Mitochondrial Dynamics in the Regulation of Nutrient Utilization and Energy Expenditure. Cell Metabolism, 2013. 17(4): p. 491–506.

58. Guido, C., et al., Mitochondrial Fission Induces Glycolytic Reprogramming in Cancer-Associated Myofibroblasts, Driving Stromal Lactate Production, and Early Tumor Growth. Oncotarget, 2012. 3(8).

59. Suh, J., et al., Mitochondrial fragmentation and donut formation enhance mitochondrial secretion to promote osteogenesis. Cell Metabolism, 2023. 35(2): p. 345–360.e7.

60. Nie, A., et al., Roles of aminoacyl-tRNA synthetases in immune regulation and immune diseases. Cell Death Dis, 2019. 10(12): p. 901.

61. Kramnik, I. and G. Beamer, Mouse models of human TB pathology: roles in the analysis of necrosis and the development of host-directed therapies. Semin Immunopathol, 2016. 38(2): p. 221–37.

62. Singh, H., et al., Meclizine ameliorates memory deficits in streptozotocin-induced experimental dementia in mice: role of nuclear pregnane X receptors. Canadian Journal of Physiology and Pharmacology, 2020. 98(6): p. 383–390.

63. Wibble, T., et al., The effects of meclizine on motion sickness revisited. Br J Clin Pharmacol, 2020. 86(8): p. 1510–1518.

64. Cohen, B. and J.M. DeJong, Meclizine and placebo in treating vertigo of vestibular origin. Relative efficacy in a double-blind study. Arch Neurol, 1972. 27(2): p. 129–35.

65. Helaine, S., et al., Host stress drives tolerance and persistence: The bane of anti-microbial therapeutics. Cell Host Microbe, 2024. 32(6): p. 852–862.

66. Sakatos, A., et al., Posttranslational modification of a histone-like protein regulates phenotypic resistance to isoniazid in mycobacteria. Sci Adv, 2018. 4(5): p. eaao1478.

67. Javid, B., et al., Mycobacterial mistranslation is necessary and sufficient for rifampicin phenotypic resistance. Proc Natl Acad Sci U S A, 2014. 111(3): p. 1132–7.

68. Rego, E.H., R.E. Audette, and E.J. Rubin, Deletion of a mycobacterial divisome factor collapses single-cell phenotypic heterogeneity. Nature, 2017. 546(7656): p. 153-157.

69. Aldridge, B.B., et al., Asymmetry and aging of mycobacterial cells lead to variable growth and antibiotic susceptibility. Science, 2012. 335(6064): p. 100- 4.

70. Stapels, D.A.C., et al., Salmonella persisters undermine host immune defenses during antibiotic treatment. Science, 2018. 362(6419): p. 1156-1160.

71. Ronneau, S., C. Michaux, and S. Helaine, Decline in nitrosative stress drives antibiotic persister regrowth during infection. Cell Host Microbe, 2023. 31(6): p. 993–1006 e6.

72. Bandyopadhyay, P., et al., S-Adenosylmethionine-responsive cystathionine beta-synthase modulates sulfur metabolism and redox balance in Mycobacterium tuberculosis. Sci Adv, 2022. 8(25): p. eabo0097.

73. Shee, S., et al., Moxifloxacin-Mediated Killing of Mycobacterium tuberculosis Involves Respiratory Downshift, Reductive Stress, and Accumulation of Reactive Oxygen Species. Antimicrob Agents Chemother, 2022. 66(9): p. e0059222.

74. Gleeson, L.E., et al., Cutting Edge: Mycobacterium tuberculosis Induces Aerobic Glycolysis in Human Alveolar Macrophages That Is Required for Control of Intracellular Bacillary Replication. J Immunol, 2016. 196(6): p. 2444–9.

75. Lachmandas, E., et al., Rewiring cellular metabolism via the AKT/mTOR pathway contributes to host defence against Mycobacterium tuberculosis in human and murine cells. Eur J Immunol, 2016. 46(11): p. 2574–2586.

76. Howard, N.C. and S.A. Khader, Immunometabolism during Mycobacterium tuberculosis Infection. Trends Microbiol, 2020. 28(10): p. 832–850.

77. Olson, G.S., et al., Type I interferon decreases macrophage energy metabolism during mycobacterial infection. Cell Rep, 2021. 35(9): p. 109195.

78. Huang, S.C., et al., Metabolic Reprogramming Mediated by the mTORC2-IRF4 Signaling Axis Is Essential for Macrophage Alternative Activation. Immunity, 2016. 45(4): p. 817–830.

79. Shi, L., et al., Infection with Mycobacterium tuberculosis induces the Warburg effect in mouse lungs. Sci Rep, 2015. 5: p. 18176.

80. Subbian, S., et al., Lesion-Specific Immune Response in Granulomas of Patients with Pulmonary Tuberculosis: A Pilot Study. PLoS One, 2015. 10(7): p. e0132249.

81. Braverman, J., et al., HIF-1alpha Is an Essential Mediator of IFN-gamma-Dependent Immunity to Mycobacterium tuberculosis. J Immunol, 2016. 197(4): p. 1287–97.

82. Marin Franco, J.L., et al., Host-Derived Lipids from Tuberculous Pleurisy Impair Macrophage Microbicidal-Associated Metabolic Activity. Cell Rep, 2020. 33(13): p. 108547.

83. Lastrucci, C., et al., Tuberculosis is associated with expansion of a motile, permissive and immunomodulatory CD16(+) monocyte population via the IL- 10/STAT3 axis. Cell Res, 2015. 25(12): p. 1333–51.

84. Beam, J.E., et al., Inflammasome-mediated glucose limitation induces antibiotic tolerance in Staphylococcus aureus. iScience, 2023. 26(10): p. 107942.

85. Beam, J.E., et al., Macrophage-Produced Peroxynitrite Induces Antibiotic Tolerance and Supersedes Intrinsic Mechanisms of Persister Formation. Infect Immun, 2021. 89(10): p. e0028621.

86. Rowe, S.E., et al., Reactive oxygen species induce antibiotic tolerance during systemic Staphylococcus aureus infection. Nat Microbiol, 2020. 5(2): p. 282–290.

87. Garaude, J., et al., Mitochondrial respiratory-chain adaptations in macrophages contribute to antibacterial host defense. Nat Immunol, 2016. 17(9): p. 1037–1045.

88. Geng, J., et al., Kinases Mst1 and Mst2 positively regulate phagocytic induction of reactive oxygen species and bactericidal activity. Nat Immunol, 2015. 16(11): p. 1142–52.

89. Wang, P., et al., Macrophage achieves self-protection against oxidative stress- induced ageing through the Mst-Nrf2 axis. Nat Commun, 2019. 10(1): p. 755.

90. Qian, Z., et al., Expression of nuclear factor, erythroid 2-like 2-mediated genes differentiates tuberculosis. Tuberculosis (Edinb), 2016. 99: p. 56–62.

91. Gatbonton-Schwager, T.N., et al., Identification of a negative feedback loop in biological oxidant formation fegulated by 4-hydroxy-2-(E)-nonenal. Redox Biol, 2014. 2: p. 755–63.

92. Ashino, T., et al., Negative feedback regulation of lipopolysaccharide-induced inducible nitric oxide synthase gene expression by heme oxygenase-1 induction in macrophages. Mol Immunol, 2008. 45(7): p. 2106–15.

93. Lewerenz, J., et al., The cystine/glutamate antiporter system x(c)(-) in health and disease: from molecular mechanisms to novel therapeutic opportunities. Antioxid Redox Signal, 2013. 18(5): p. 522–55.

94. Vilcheze, C. and W.R. Jacobs, Jr., The promises and limitations of N- acetylcysteine as a potentiator of first-line and second-line tuberculosis drugs. Antimicrob Agents Chemother, 2023. 65(5).

95. Zhou, J., et al., Activation of Nrf2 modulates protective immunity against Mycobacterium tuberculosis infection in THP1-derived macrophages. Free Radic Biol Med, 2022. 193(Pt 1): p. 177–189.

96. Beam, J.E., et al., The Use of Acute Immunosuppressive Therapy to Improve Antibiotic Efficacy against Intracellular Staphylococcus aureus. Microbiol Spectr, 2022. 10(3): p. e0085822.

97. Finkel, T., Signal transduction by reactive oxygen species. J Cell Biol, 2011. 194(1): p. 7–15.

98. Sinenko, S.A., et al., Physiological Signaling Functions of Reactive Oxygen Species in Stem Cells: From Flies to Man. Front Cell Dev Biol, 2021. 9: p. 714370.

99. Bonnet, S., et al., A mitochondria-K+ channel axis is suppressed in cancer and its normalization promotes apoptosis and inhibits cancer growth. Cancer Cell, 2007. 11(1): p. 37–51.

100. Chen, Q., et al., Modulation of electron transport protects cardiac mitochondria and decreases myocardial injury during ischemia and reperfusion. Am J Physiol Cell Physiol, 2007. 292(1): p. C137–47.

101. Piantadosi, C.A. and J. Zhang, Mitochondrial generation of reactive oxygen species after brain ischemia in the rat. Stroke, 1996. 27(2): p. 327–31; discussion 332.

102. Padmapriydarsini, C., et al., Randomized Trial of Metformin With Anti- Tuberculosis Drugs for Early Sputum Conversion in Adults With Pulmonary Tuberculosis. Clin Infect Dis, 2022. 75(3): p. 425–434.

103. Hong, C.T., K.Y. Chau, and A.H. Schapira, Meclizine-induced enhanced glycolysis is neuroprotective in Parkinson disease cell models. Sci Rep, 2016. 6: p. 25344.

104. Gohil, V.M., et al., Meclizine is neuroprotective in models of Huntington’s disease. Hum Mol Genet, 2011. 20(2): p. 294–300.

105. Lione, A. and A.R. Scialli, The developmental toxicity of the H1 histamine antagonists. Reprod Toxicol, 1996. 10(4): p. 247–55.

106. Giurgea, M. and J. Puigdevall, Experimental teratology with Meclozine. Med Pharmacol Exp Int J Exp Med, 1966. 15(4): p. 375–88.

107. Chandra, P., S.J. Grigsby, and J.A. Philips, Immune evasion and provocation by Mycobacterium tuberculosis. Nat Rev Microbiol, 2022. 20(12): p. 750–766.

108. Gern, B.H., et al., TGFbeta restricts expansion, survival, and function of T cells within the tuberculous granuloma. Cell Host Microbe, 2021. 29(4): p. 594–606 e6.

109. Gautam, U.S., et al., In vivo inhibition of tryptophan catabolism reorganizes the tuberculoma and augments immune-mediated control of Mycobacterium tuberculosis. Proc Natl Acad Sci U S A, 2018. 115(1): p. E62–E71.

110. Cronan, M.R., et al., Macrophage Epithelial Reprogramming Underlies Mycobacterial Granuloma Formation and Promotes Infection. Immunity, 2016. 45(4): p. 861–876.

111. Guler, R., et al., Targeting Molecular Inflammatory Pathways in Granuloma as Host-Directed Therapies for Tuberculosis. Front Immunol, 2021. 12: p. 733853.

112. Putri, G.H., et al., Analysing high-throughput sequencing data in Python with HTSeq 2.0. Bioinformatics, 2022. 38(10): p. 2943–2945.

113. Love, M.I., W. Huber, and S. Anders, Moderated estimation of fold change and dispersion for RNA-seq data with DESeq2. Genome Biology, 2014. 15(12): p. 550.

114. Liberzon, A., et al., Molecular signatures database (MSigDB) 3.0. Bioinformatics, 2011. 27(12): p. 1739–1740.

115. Martinez, F.O., et al., Transcriptional profiling of the human monocyte-to-macrophage differentiation and polarization: new molecules and patterns of gene expression. J Immunol, 2006. 177(10): p. 7303–11.

116. Fang, Z., X. Liu, and G. Peltz, GSEApy: a comprehensive package for performing gene set enrichment analysis in Python. Bioinformatics, 2023. 39(1).

117. Robinson, M.D., D.J. McCarthy, and G.K. Smyth, edgeR: a Bioconductor package for differential expression analysis of digital gene expression data. Bioinformatics, 2010. 26(1): p. 139–40.

118. Khan, M.Z., et al., Protein kinase G confers survival advantage to Mycobacterium tuberculosis during latency-like conditions. J Biol Chem, 2017. 292(39): p. 16093–16108.

119. Jain, R., et al., Enhanced and enduring protection against tuberculosis by recombinant BCG-Ag85C and its association with modulation of cytokine profile in lung. PLoS One, 2008. 3(12): p. e3869.

120. Dutta, N.K., M.L. Pinn, and P.C. Karakousis, Reduced emergence of isoniazid resistance with concurrent use of thioridazine against acute murine tuberculosis. Antimicrob Agents Chemother, 2014. 58(7): p. 4048–53.

