## Supplementary data for "Bioenergetic reprogramming of macrophages reduces drug tolerance in *Mycobacterium tuberculosis*"

**Supplementary materials for “Bioenergetic reprogramming of macrophages reduces drug tolerance in *Mycobacterium tuberculosis*”**

Fig. S1. Gating strategy to flow-sort macrophages with $E_{MSH}$-reduced or $E_{MSH}$–oxidized *Mtb.*

Fig. S2. *Mtb* subpopulations inside macrophages show differential refractoriness to multiple anti-TB drugs.

Fig S3. Macrophage subpopulations upon *Mtb* infection show heterogenous transcriptomic response

Fig S4. Gene ontology and transcription factor motif enrichment analysis.

Fig S5. Pathway enrichment of genes upregulated in macrophages enriched in $E_{MSH}$-oxidized *Mtb* compared to macrophages harboring $E_{MSH}$-reduced *Mtb* using Enrichr.

Fig. S6. Reinfection of bystander macrophages causes massive oxidative stress in *Mtb.*

Fig. S7. Mitochondrial ROS is produced through RET.

Fig. S8. siRNA mediated knockdown of Nrf2.

Fig. S9. Sorafenib (SFN) treatment is not cytotoxic to macrophages and shows a high MIC against *Mtb*.

Fig. S10. Macrophage metabolism affects antibiotic tolerance of intracellular *Mtb.*

Fig. S11. MEC is not cytotoxic to macrophages and does not affect *Mtb* in-vitro.

Fig. S12. Transcriptional profiling of infected macrophages treated with MEC.

Fig. S13. *Mtb*-infection decreases oxidative phosphorylation in BMDMs.

**Other Supplementary Material for this manuscript includes the following (provided as MS Excel files):**

Table S1- List of differentially expressed genes between the infected macrophage subpopulations- bystanders, oxidized and reduced compared to uninfected BMDMs.

Table S2- List of differentially expressed genes between the reduced and oxidized subpopulations.

Table S3- List of genes used for overlap analysis of DEGs from macrophage subpopulations compared to uninfected from this study with infected BMDMs 24h p.i. from *Andreu et al., Sci Rep (2017).*

Table S4- List of genes used for overlap analysis of DEGs between ‘Reduced’ and ‘Oxidized’ subpopulations from this study with *Mtb*-infected alveolar and interstitial macrophages (AMs and IMs, respectively) from *Huang et al., JEM (2018).*

Table S5- List of genes used for overlap analysis of DEGs between ‘Reduced’ and ‘Oxidized’ subpopulations from this study with *Mtb*-infected murine alveolar macrophages isolated from BAL fluid 10d p.i. from *Rothchild et al., Sci Immunol (2019).*

Table S6- List of genes used for overlap analysis of DEGs between ‘Reduced’ and ‘Oxidized’ subpopulations from this study with M1 and M2 macrophage gene signatures taken from GEO (accession number- GSE5099)

Table S7- List of differentially expressed genes between infected macrophages treated with 20 µM MEC or 0.2 % DMSO (solvent control). Genes downregulated upon MEC treatment are highlighted in blue, and upregulated are highlighted in red.

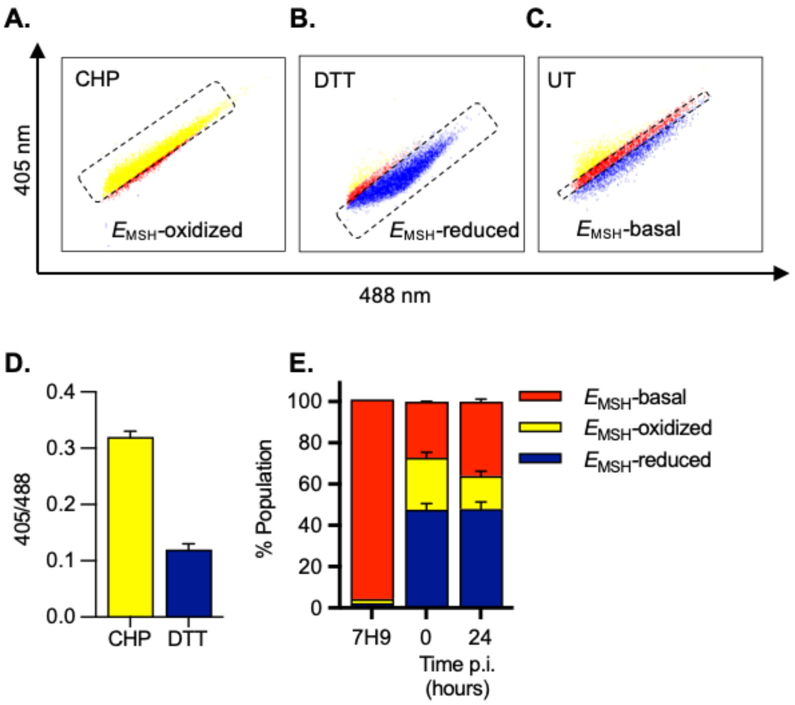


**Fig. S1. Gating strategy to flow-sort macrophages with $E_{MSH}$-reduced or $E_{MSH}$–oxidized *Mtb.*** Cumene hydroperoxide (CHP) and dithiothriotol (DTT) were used independently to gate the $E_{MSH}$-oxidized and –reduced subpopulations, respectively. Dot plots represent cells that were majorly oxidized or reduced upon treatment with **A.** 10 mM CHP (shown in yellow) or **B.**100 mM DTT (shown in blue), respectively; **C.** the remaining subpopulation was gated as $E_{MSH}$-basal. **D.** Ratiometric response of biosensor to CHP and DTT at the two excitation wavelengths- 405 nm and 488 nm with emission recorded at 510 nm; **E.** Bar plots showing redox heterogeneity of *Mtb* grown in vitro in 7H9 medium and in macrophages at 0 and 24 h post-infection (p.i.). Data are expressed as mean ± S.D. of three independent experiments.

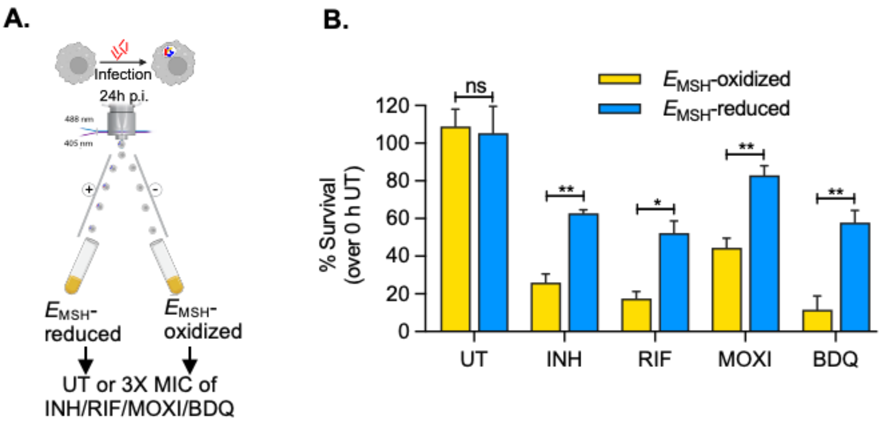


**Fig. S2. *Mtb* subpopulations inside macrophages show differential refractoriness to multiple anti-TB drugs**. **A.** Schematic showing the experimental strategy. Infected BMDMs predominantly harboring either $E_{MSH}$-reduced or $E_{MSH}$-oxidized *Mtb* were flow-sorted 24 hours p.i. and were either left untreated or further treated with 3X MIC of different anti-TB drugs- isoniazid (INH; MIC-0.125 ug/ml), rifampicin (RIF; 0.125 ug/ml), moxifloxacin (MOXI, MIC- 0.25 ug/ml), or bedaquiline (BDQ, MIC- 0.0625 ug/ml) for additional 48 hours. **B.** Bar plot showing the percentage survival of intracellular *Mtb* post 48 h treatment. Percentage survival was calculated by normalizing the CFU in antibiotic-treated cells at 48 hours against untreated cells at 0 hours. Data are expressed as mean ± S.D. of three independent experiments. p-value was determined using an unpaired t-test with Welch's correction. ns- non-significant, *$p$<0.05, **$p$<0.01, ***$p$<0.001, ****$p$<0.0001.

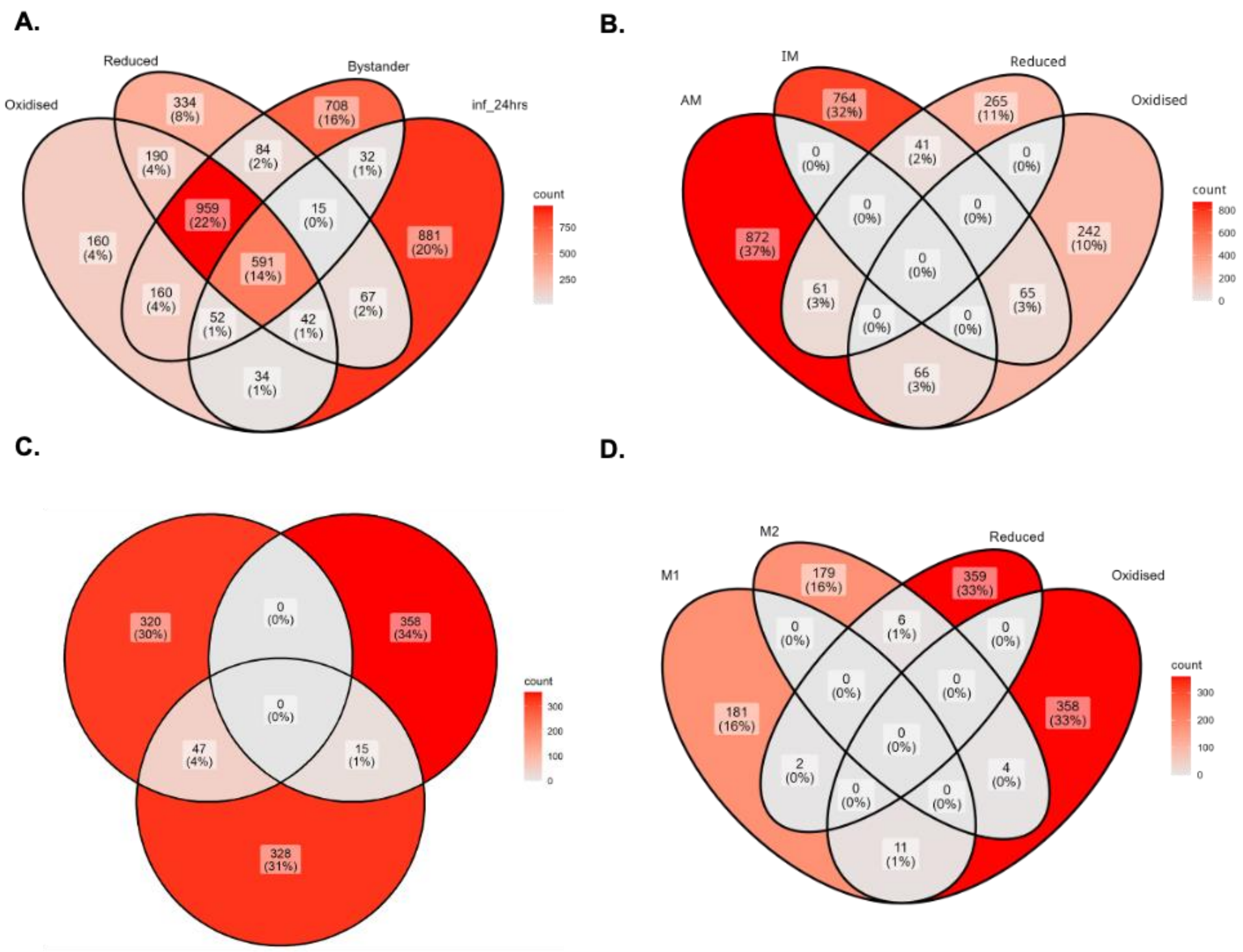


**Fig. S3. Macrophage subpopulations upon *Mtb* infection show heterogenous transcriptomic response. A.** Overlap analysis between DEGs in the mentioned subpopulations compared to naïve uninfected macrophages from this study with those found in published literature (infected vs. uninfected at 24 h p.i. in *Andreu et al., Sci rep (2017))*; overlap analysis between reduced and oxidized macrophage subpopulations from this study with **B.** *Mtb*-infected AMs and IMs from *Huang et al., JEM (2018);* **C.** Infected AMs isolated from BAL fluid after 10 days of infection from *Rothchild et al., Sci Immunol. (2019)*; **D.** M1 and M2 macrophage populations obtained from GEO database (accession number GSE5099).

**A.**

| Rank | TF | Set name | Set size | Overlapping Genes | FET p-value | FDR | Odds ratio |
|---|---|---|---|---|---|---|---|
| 1 | NFE2L2 | NFE2L2_20460467_CHIPSEQ_MEF_MOUSE | 942 | 48 | 5.70E-12 | 8.76E-10 | 3.502 |
| 2 | GABPA | GABPA_20460467_CHIPSEQ_MEF_MOUSE | 942 | 48 | 5.70E-12 | 8.76E-10 | 3.502 |
| 3 | MTF2 | MTF2_20144788_CHIPSEQ_MEF_MOUSE | 2645 | 88 | 7.10E-10 | 7.277E-8 | 2.245 |
| 4 | ZNF217 | ZNF217_24962896_CHIPSEQ_MEF_HUMAN | 1413 | 54 | 8.44E-9 | 6.487E-7 | 2.592 |
| 5 | MYB | MYB_21317192_CHIPSEQ_ERMYB_MOUSE | 837 | 38 | 1.32E-8 | 8.13E6-7 | 3.102 |
| 6 | MITF | MITF_21258399_CHIPSEQ_MELANOMA_HUMAN | 5140 | 136 | 8.64E-8 | 4.42E-6 | 1.773 |
| 7 | WT1 | WT1_25993318_CHIPSEQ_PODOVYTE_HUMAN | 3079 | 90 | 1.19E-7 | 5.24E-6 | 1.964 |
| 8 | NR1I2 | NR1I2_20693526_CHIPSEQ_LIVER_MOUSE | 715 | 32 | 2.09E-7 | 8.02E-6 | 3.056 |
| 9 | ESR1 | ESR1_22446102_CHIPSEQ_UTERI_MOUSE | 1320 | 47 | 4.32E-7 | 1.48E-5 | 2.408 |
| 10 | FOXM1 | FOXM1_26456572_CHIPSEQ_MCF7_HUMAN | 1828 | 58 | 1.02E-6 | 3.15E-5 | 2.137 |

**B.**

| Enrichment FDR | nGenes | Pathway Genes | Fold enrichment | Pathways |
|---|---|---|---|---|
| 3.0E-24 | 41 | 365 | 9.1 | BACH2 KO Mouse GSE54050 **UP** |
| 1.0E-05 | 14 | 187 | 6.0 | NFE2L2 KO Mouse GSE18344 **DOWN** |

**C.**

| Enrichment FDR | nGenes | Pathway Genes | Fold enrichment | Pathways |
|---|---|---|---|---|
| 2.2E-02 | 3 | 14 | 17.3 | WP454 OSTEOCLAST SIGNALING |
| 1.2E-03 | 5 | 28 | 14.4 | WP412 OXIDATIVE STRESS RESPONSE |
| 4.2E-02 | 3 | 19 | 12.8 | WP164 GLUTATHIONE METABOLISM |
| 4.3E-02 | 3 | 22 | 11 | WP336 FATTY ACID BIOSYNTHESIS |
| 2.3E-04 | 9 | 92 | 7.9 | WP4466 OXIDATIVE STRESS AND REDOX PATHWAY |
| 4.3E-02 | 8 | 185 | 3.5 | WP85 FOCAL ADHESION |

**Fig. S4. Gene ontology and transcription factor motif enrichment analysis. A.** ChEA3-based enrichment analysis of transcription factor(TF) motif in the promoter regions of DEGs between redox diverse subpopulations. The Nrf2 binding motif is among the top TF motifs in the promoter regions of the reduced population gene signature; **B.** TF perturbation followed by expression changes from available datasets compared to genes induced in macrophages with $E_{MSH}$-reduced *Mtb;* **C.** ShinyGO-based pathway enrichment of genes upregulated in macrophages with $E_{MSH}$-reduced *Mtb*.

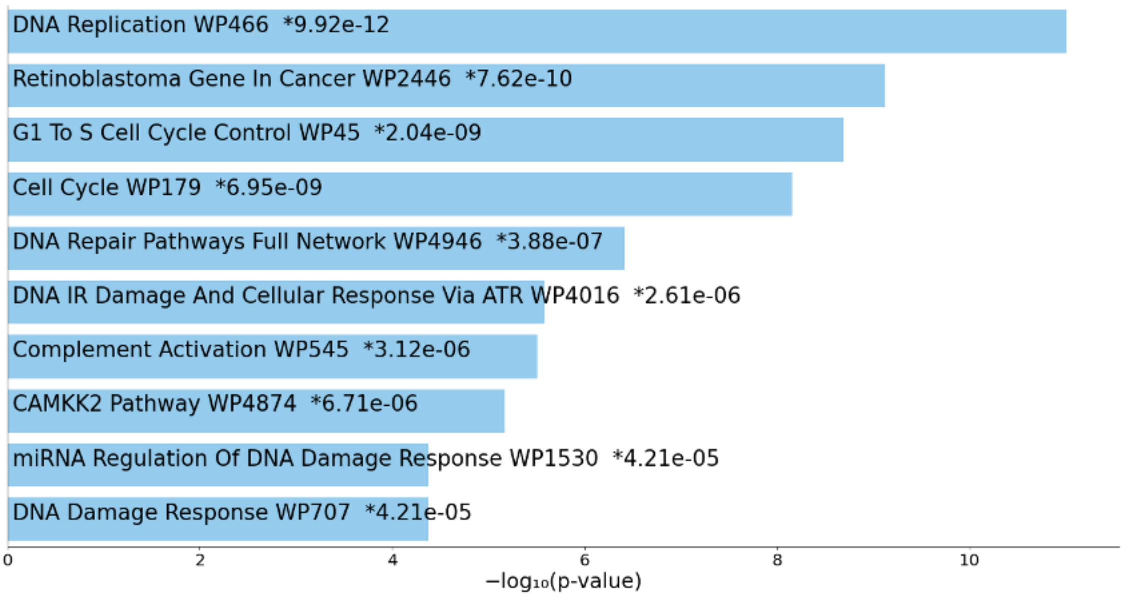


**Fig. S5. Pathway enrichment of genes upregulated in macrophages enriched in $E_{MSH}$-oxidized *Mtb* compared to macrophages harboring $E_{MSH}$-reduced *Mtb* using Enrichr.**

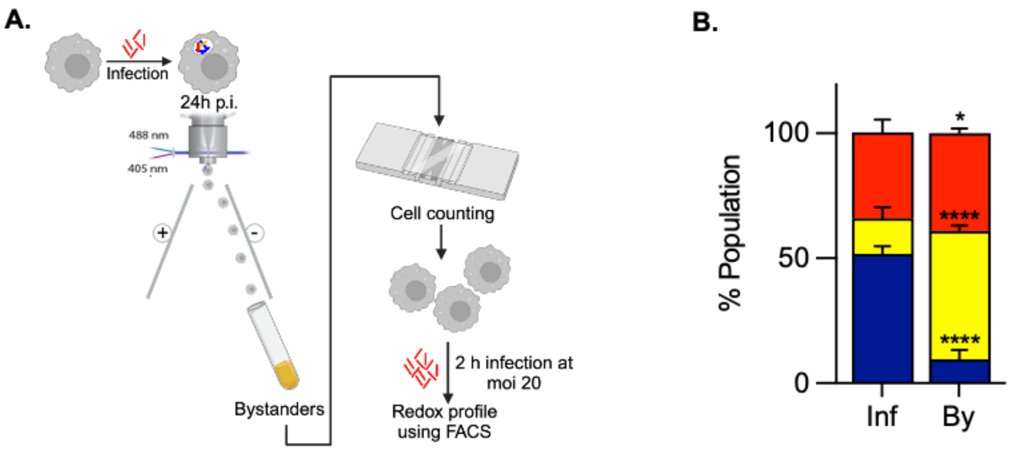


**Fig. S6. Reinfection of bystander macrophages causes massive oxidative stress in *Mtb.* A.** Schematic showing the experimental plan. BMDMs were initially infected with *Mtb*-roGFP2, and the bystander macrophages were flow sorted. These cells were reinfected with *Mtb*-roGFP2 for 2 hours and the redox diversity was assessed using FACS. **B.** Bar plot showing redox diversity in BMDMs post first round of infection ("Inf") and in bystander macrophages post reinfection ("By"). Data are expressed as mean ± S.D. of three biological replicates done at least in triplicates. p-value determined using an unpaired t-test with Welch's correction. ns- non-significant, *$p$<0.05, **$p$<0.01, ***$p$<0.001, ****$p$<0.0001

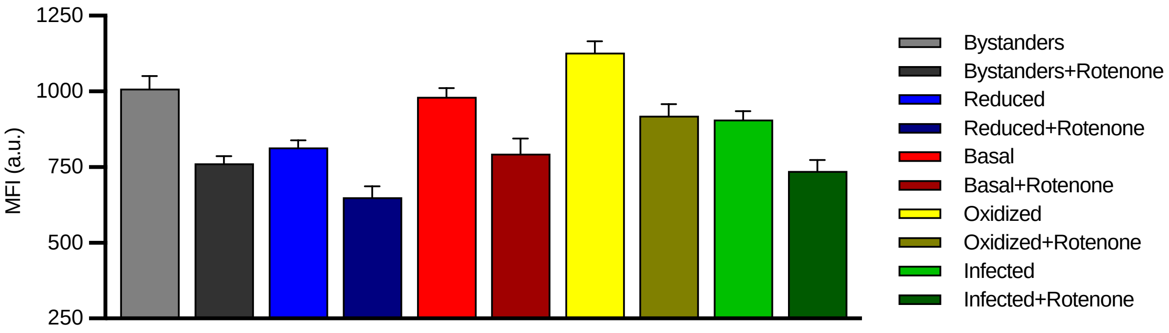


**Fig. S7. Mitochondrial ROS is produced through RET.** Bar plot showing median fluorescence intensity (MFI) of MitoSOX in different macrophage subpopulations. *Mtb*-roGFP2 infected BMDMs were treated with 10 µM rotenone for 1 h before staining. Data are expressed as mean ± S.D. representative of two independent experiments.

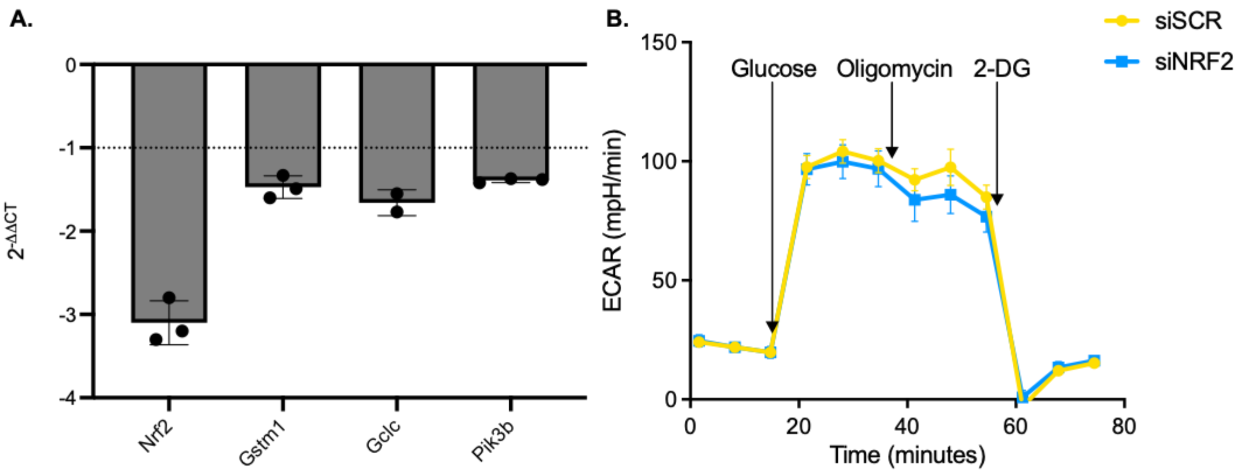


**Fig. S8. siRNA mediated knockdown of Nrf2. A.** qRT-PCR for Nrf2 and Nrf2-regulon genes. The bar plot shows gene expression in BMDMs transfected with siRNA against Nrf2 (siNRF2) compared to scrambled siRNA (siSCR)-transfected BMDMs; **B.** Glycolytic function test of infected BMDMs at 24 h p.i. upon Nrf2 KD.

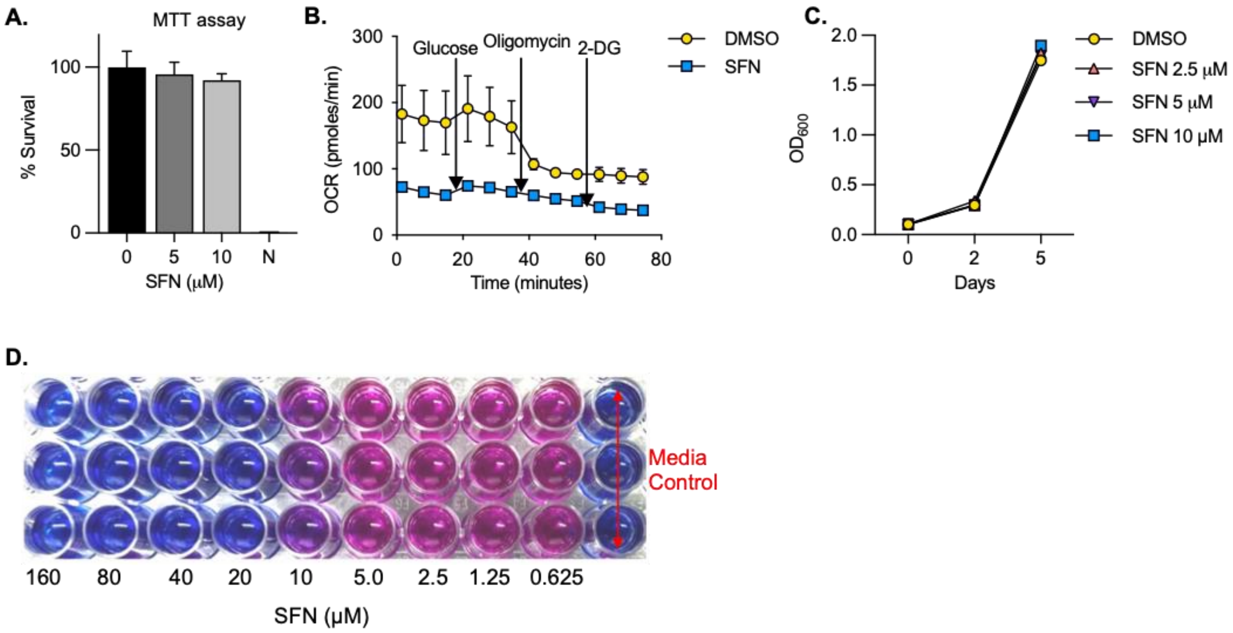


**Fig. S9. Sorafenib (SFN) treatment is not cytotoxic to macrophages and shows a high MIC against *Mtb*. A.** MTT assay to assess the survival of macrophages upon treatment with different concentrations of SFN for 24 h; **B.** Oxygen consumption rate upon treatment with 5 µM SFN in *Mtb*-infected BMDMs at 24 h p.i.; **C.** In-vitro survival of *Mtb* upon treatment with varying concentrations of SFN assessed by measuring optical density at 600 nm; **D.** MIC of SFN against *Mtb* determined by Resazurin Microtiter Assay (REMA). MIC=20 µM.

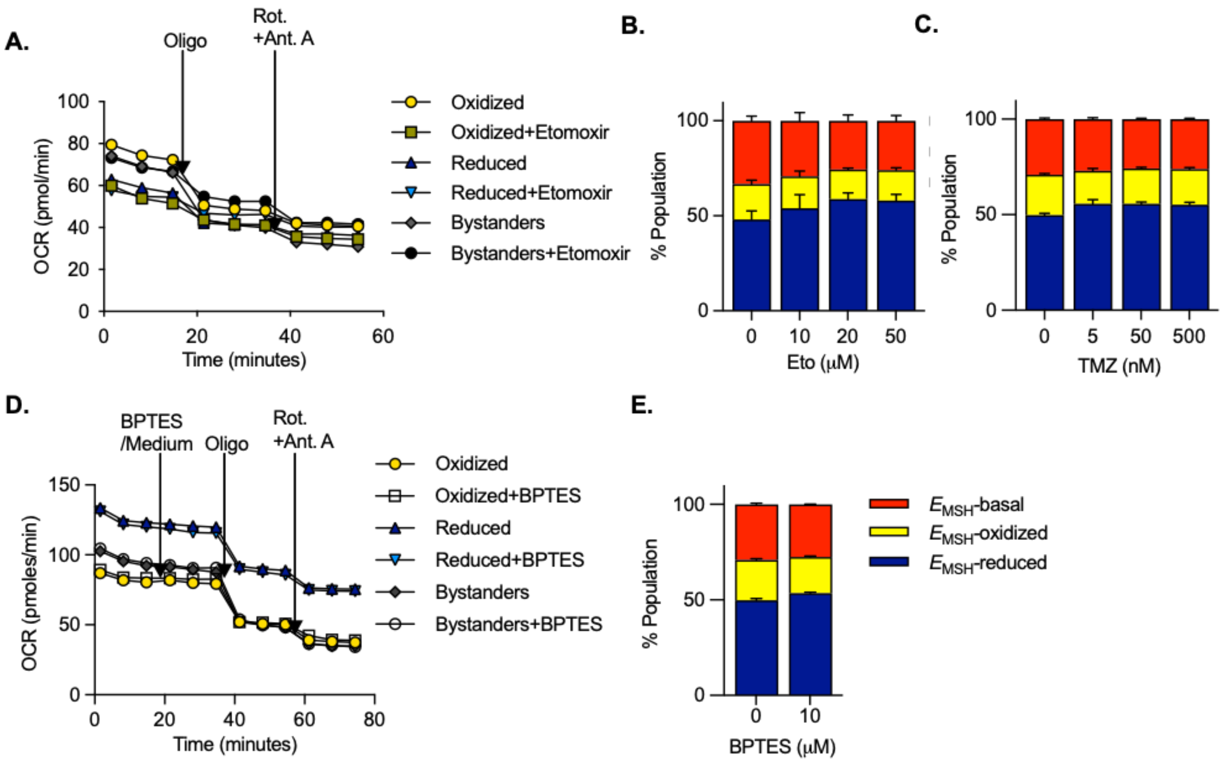


**Fig. S10. Macrophage metabolism affects antibiotic tolerance of intracellular *Mtb*. A.** Modified mitostress test was done to determine the contribution of fatty acid oxidation towards oxidative phosphorylation in *Mtb*-infected BMDM subpopulations. Cells were treated with 20µM etomoxir (Eto) for 24 h p.i. and subjected to the assay; **B.** Redox profile of intracellular *Mtb* in macrophages treated with either 10, 20 or 50µM Eto for 24h p.i.; **C.** Redox profile of intracellular *Mtb* in macrophages treated with either 5, 50 or 500nM trimetazidine (TMZ) 24h p.i.; **D.** Mitostress test to determine the contribution of glutamine towards oxidative phosphorylation in *Mtb*-infected macrophage subpopulations at 24h p.i.. At the indicated time point, either 10µM BPTES or culture medium (control) was added to the cells, and OCR was measured; **E.** Redox profile of intracellular *Mtb* at 24h p.i. in macrophages upon treatment with 10µM BPTES**.** Data are expressed as mean ± S.D. representative of two independent experiments. All changes are non-significant, determined by an unpaired t-test with Welch's correction.

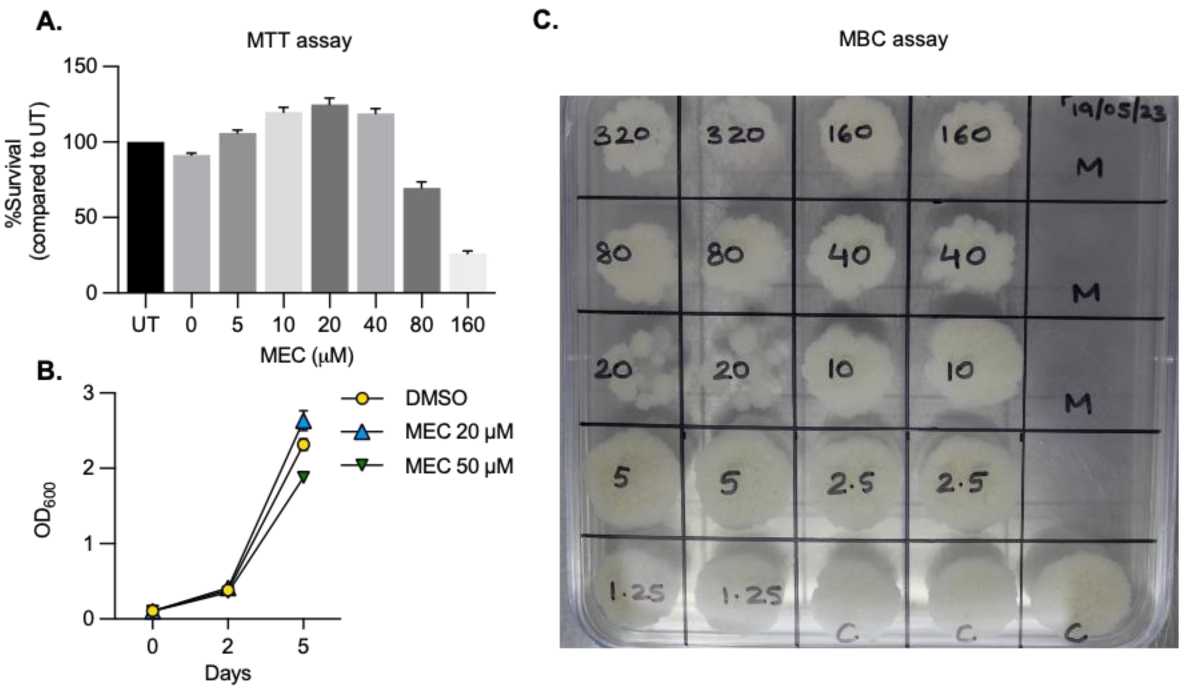


**Fig. S11. MEC is not cytotoxic to macrophages and does not affect *Mtb* in-vitro.** **A.** MTT (3-(4, 5-dimethylthiazolyl-2)-2, 5-diphenyltetrazolium bromide) assay to assess the survival of macrophages upon 24 h treatment with MEC at the indicated concentrations. Survival was normalized to untreated macrophages; **B.** *Mtb* growth in vitro was assessed by measuring optical density at 600 nm; **C.** Minimum bactericidal concentration (MBC) was assessed by plating the culture from the REMA plate on solid medium (7H11 agar + OADC). MBC- >320µM.

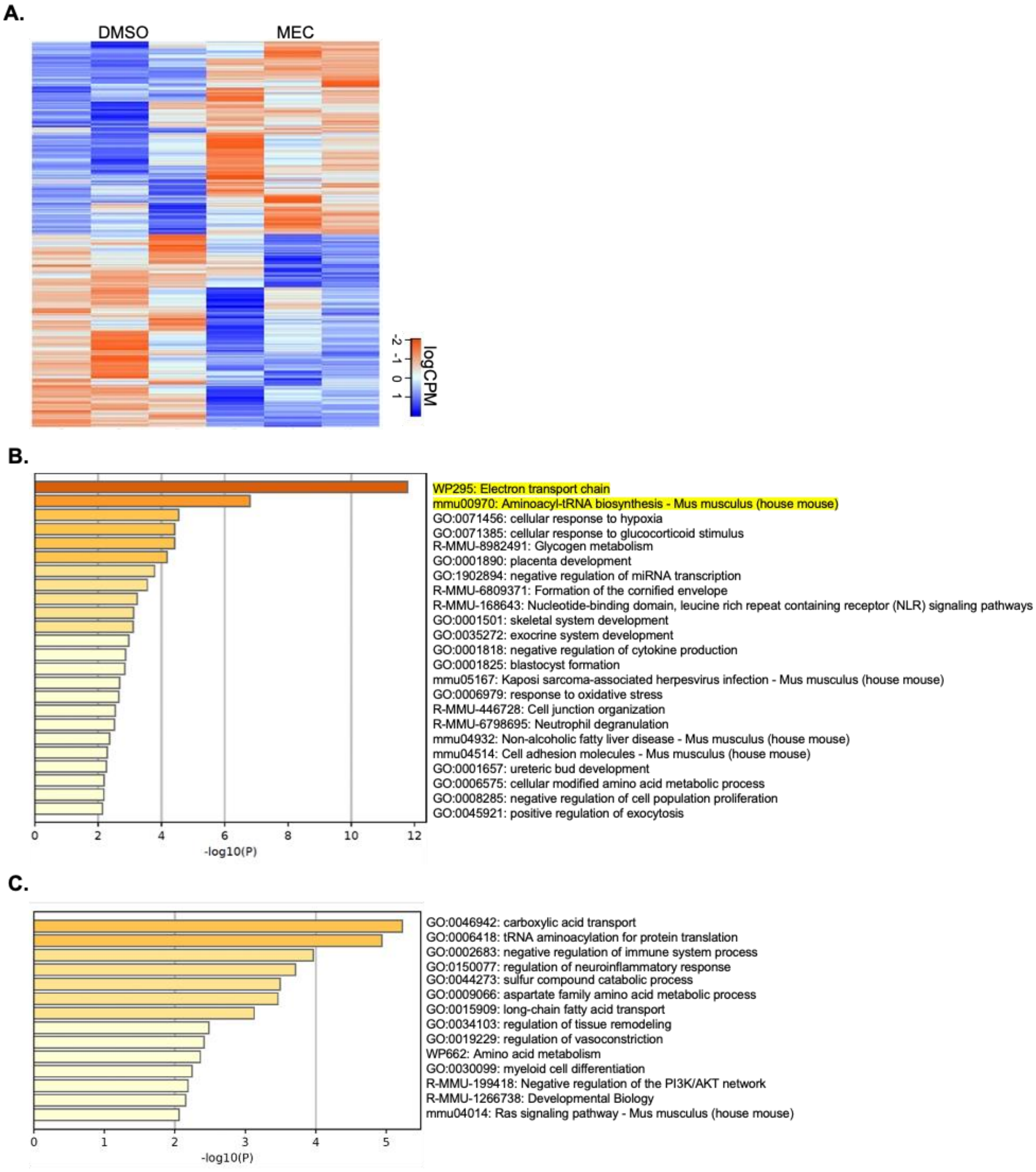


**Fig. S12. Transcriptional profiling of infected macrophages treated with MEC. A.** Heatmap showing differentially expressed genes between macrophages treated either with 20µM meclizine or equivalent 0.2% DMSO control**.** GFP+ infected macrophages were flow-sorted, and RNA sequencing was performed. For the heatmap, basemean is >10, and FDR is <0.1**; B.** Major pathways downregulated upon treatment with MEC; **C.** major pathways upregulated upon treatment with MEC compared to DMSO control. Metascape pathway enrichment tool was used for pathway enrichment analysis.

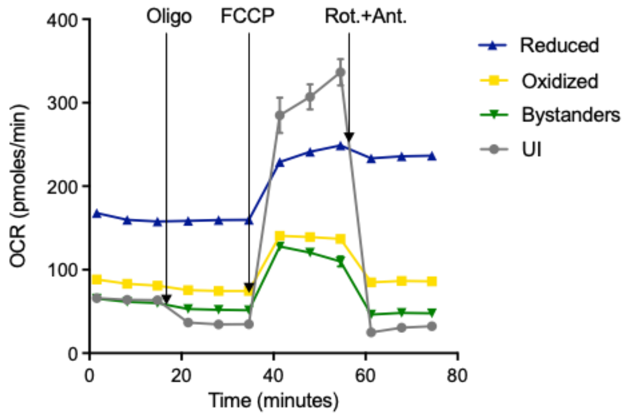


**Fig. S13. *Mtb*-infection decreases oxidative phosphorylation in BMDMs.** UI represents uninfected macrophages while reduced, oxidized, and bystanders are the macrophage subpopulations seen 24 h p.i. At the indicated time points, 2 µM oligomycin (oligo), 1 µM FCCP, 2 µM rotenone and 4 µM antimycin A were added.
